# A new twist on an old idea: The *Posidonia oceanica* Large PSI-LHCI-LHCII supercomplex provides the structural basis of photosystem I spectral diversification in higher plants

**DOI:** 10.64898/2026.04.14.718421

**Authors:** Quentin Charras-Ferroussier, Tahani Alsenani, Ashraf Al-Amoudi, Marina I. Siponen, Emmanuel Heilmann, Andreas Naschberger, Colette Jungas

**Affiliations:** Science for Life Laboratory, School of Engineering Sciences in Chemistry, Biotechnology and Health, KTH Royal Institute of Technology, KTH University, Stockholm, Sweden; King Abdullah University of Science and Technology (KAUST), Biomedical Sciences (BioMed) Division, 23955 Thuwal, Saudi Arabia; King Abdullah University of Science and Technology (KAUST), Electron Microscopy Lab, Imaging and Characterization Core Labs, Thuwal 23955, Saudi Arabia; Aix Marseille University, CEA, CNRS, BIAM, Team Proteintec, 13115 Saint Paul-Lez-Durance, France; Aix Marseille University, CEA, CNRS, BIAM, Team LGBP, 13115 Saint Paul-Lez-Durance, France

## Abstract

Higher plant Photosystem I (PSI) can capture low-energy photons and convert them into chemical energy through special far-red absorbing chlorophylls, the red-forms. Across higher-plant evolution, red-form spectral range has shifted to match differences in far-red availability across various habitats, despite broad conservation of the antenna system and canonical red-form architecture. Here, we report the 1.9 Å cryo-EM structure of a large and blue shifted PSI light-harvesting complex I and II supercomplex (L-PSI-LHCI-LHCII) from the Mediterranean seagrass *Posidonia oceanica*, a marine angiosperm adapted to blue-enriched underwater light depleted in far-red wavelengths. Although the PSI core remains conserved, the antenna system was selectively remodeled through incorporation of an additional Lhca1–Lhca4 heterodimer at a previously undescribed binding site, together with pigment remodeling in the LHCI and the LHCII that enhanced blue–cyan light harvesting. Comparative structural analyses identified recurrent amino-acid sites within the far-red domains that influence red-forms geometry and whose residue composition diverges strongly among blue-shifted and red-shifted plant species. These findings establish a common structural basis underlying the spectral diversification of PSI toward contrasting light environments, accounting for both seagrass adaptation to the seawater light regime and the broad range of far-red absorption in higher plants.

## INTRODUCTION

In plant photosynthesis, Photosystem I (PSI) and Photosystem II (PSII) absorb sunlight through chlorophyll *a*/*b*-binding proteins, the light-harvesting complexes (LHCs). LHCII, primarily associated with PSII, is comprised of monomeric and trimeric complexes assembled with the PSII core into PSII–LHCII supercomplexes which are largely conserved across green plant lineages (Su et al., 2017; Shen et al., 2019; Kouřil et al., 2020). In addition, a mobile pool of trimeric LHCII can be reversibly recruited to PSI, acting as a shared antenna system that balances excitation between the two photosystems under fluctuating light environments (Wientjes et al., 2013; Goldschmidt-Clermont and Bassi, 2015; Albanese et al., 2016). On the other hand LHCI, the peripheral antenna of PSI, is constitutively bound to the PSI core and has undergone a marked remodelling during evolution with a noticeable downsizing in its cross-section.

While early-branching lineages such as chlorophytes and bryophytes express large PSI supercomplexes, with antenna systems expanding up to dozen of subunits (Huang et al., 2021; Zhang et al., 2023), higher-plant LHCI shrank to a four-subunit antenna belt arranged in two heterodimers (Lhca1/4 and Lhca2/3) (Wang et al., 2021; Wu et al., 2023). This antenna reduction occurred alongside the evolution of low-energy chlorophylls, the red forms, located in LHCI and thereby extending the functional spectral range of PSI into far-red wavelengths which are abundant in the habitats of land plants. Together, the combination of these traits likely contributed to the ecological success of land plants by limiting PSI overexcitation under high irradiance in open land environments (Alboresi et al., 2009; Kono et al., 2017), while enhancing light harvesting, especially in plant dense communities, where upper canopy leaves filter sunlight into a dimmer far-red enriched spectrum (Zhen et al., 2022).

Red forms arose from specific excitonic interactions in the Chl *a*603–*a*609 pair located in Lhca3 and Lhca4 (Wientjes and Croce, 2011), where a His→Asn substitution at the axial ligand of Chl *a*603 favored the geometry and energy states enabling far-red absorption (Romero et al., 2009; Wientjes et al., 2012). This change shifted the low-temperature fluorescence emission maxima from ∼683 nm in the Lhca’s of green algae retaining the His ligand to ∼715 nm in the antenna containing the Asn substitution (Drop et al., 2011; Su et al., 2019). Yet, PSI spectral diversification continued beyond this transition, with emission maximum reaching ∼725 nm in moss (Gorski et al., 2022), 735 nm in most vascular plants (Stolárik et al., 2018; Bos et al., 2023; Colpo et al., 2023) and up to ∼745 nm in some shade-adapted angiosperms (flowering plants) (Li et al., 2024; Zer et al., 2025). This spectral diversification occurred in land plants despite conservation of the canonical LHCI architecture and the retention of Asn in Lhca3 and Lhca4, indicating that far-red energetics can be extensively remodeled without altering the canonical red-form architecture. Seagrasses are an interesting case to illustrate this, as they thrive in environments depleted in red and far-red light. In these plants, PSI fluorescence emission shifted back toward ∼720 nm despite retention of the Asn ligand (Ferroussier et al., 2025).

The endemic Mediterranean seagrass *Posidonia oceanica* (Alismatales), a marine angiosperm, is a key ecosystem engineer, acting as a major long-term blue-carbon sink (Díaz-Almela and Duarte, 2008; Pergent-Martini et al., 2021). Its meadows extend from the water surface up to a 40 m depth, across which irradiance decreases up to 50-fold and shifts toward blue-cyan wavelengths as red and far-red wavelengths progressively disappear. *P. oceanica* adapted to this environment through a blue-shifted PSI marked by strongly attenuated red forms and Chl *b-enrichment* (Ferroussier et al., 2025). The presence of a formyl (–CHO) group in Chl *b* shifts its absorption spectrum relative to Chl *a*, resulting in stronger absorption in the blue-cyan region around 453 nm, dominant in seawater, compared with the 430 nm maximum of Chl *a*. The *P. oceanica* PSI further associates with a canonically bound phosphorylated LHCII trimer and an additional Lhca1–Lhca4 heterodimer attached through a previously undescribed binding site, forming a highly abundant large PSI–LHCI–LHCII supercomplex (L-PSI-LHCII). This organization appears to have arisen repeatedly during seagrass evolution despite multiple independent transitions to seawater (Wissler et al., 2011; Chen et al., 2022; Ferroussier et al., 2025), and is notably conserved in *Zostera marina* and *Halophila stipulacea*, two of the most widespread seagrasses in the Northern Hemisphere (Scheibling et al., 2018; Yu et al., 2023). Despite this expanded antenna system, PSI maintains highly efficient excitation-energy trapping while exhibiting strongly reduced far-red emission (Ferroussier et al., 2025).

Here, we report the 1.9 Å cryo-electron microscopy structure of the *P. oceanica* L-PSI-LHCII. The high resolution of this structure allowed us to examine how antenna architecture and photosynthetic pigment composition were precisely remodeled under the seawater light regime. This structure provides a unique opportunity to compare a blue-shifted higher-plant PSI that still retains canonical red forms with its red-shifted counterparts in land plants. Together, this comparative structural analysis shows that *P. oceanica* adapted PSI to a blue– cyan-enriched marine niche through targeted remodeling of the LHC antenna system rather than extensive architectural innovation. The structure identifies the basis for additional Lhca1– Lhca4 recruitment, pigment substitutions associated with blue–cyan light harvesting, and recurrent amino-acid determinants that reshape the red-form environment across blue- and red-shifted species. These findings further reveal distinct evolutionary trajectories in Lhca3 and Lhca4 and establish red forms as a tunable system underlying the spectral plasticity of higher-plant PSI.

## RESULTS

### The structure of the L-PSI-LHCII complex

The single-particle cryo-EM structure of L-PSI-LHCII was determined utilizing 182,928 particles at a resolution of 1.92 Å for the consensus map, with local refinement up to 1.87 Å for PSI-LHCI (**Supplemental Figure 1, 2 and Supplemental Table 1**). The high-quality map facilitated the precise identification and modeling of 19 subunits (**Figure 1A and 1B**), including PsaN (absent or incompletely resolved in previous *A. thaliana* (Wu et al., 2023) and maize structures (Pan et al., 2018)) as well as 186 Chl *a*, 45 Chl *b*, 56 carotenoids (**Figure 1C**), 85 aliphatic chains, and 1007 structured water molecules. The additional Lhca1-Lhca4 dimer (Lhca1²-Lhca4²) is fully resolved and adopts the canonical architecture, with each subunit coordinating 21 Chl *a,* 8 Chl *b,* and 7 carotenoids.

**Figure 1:**
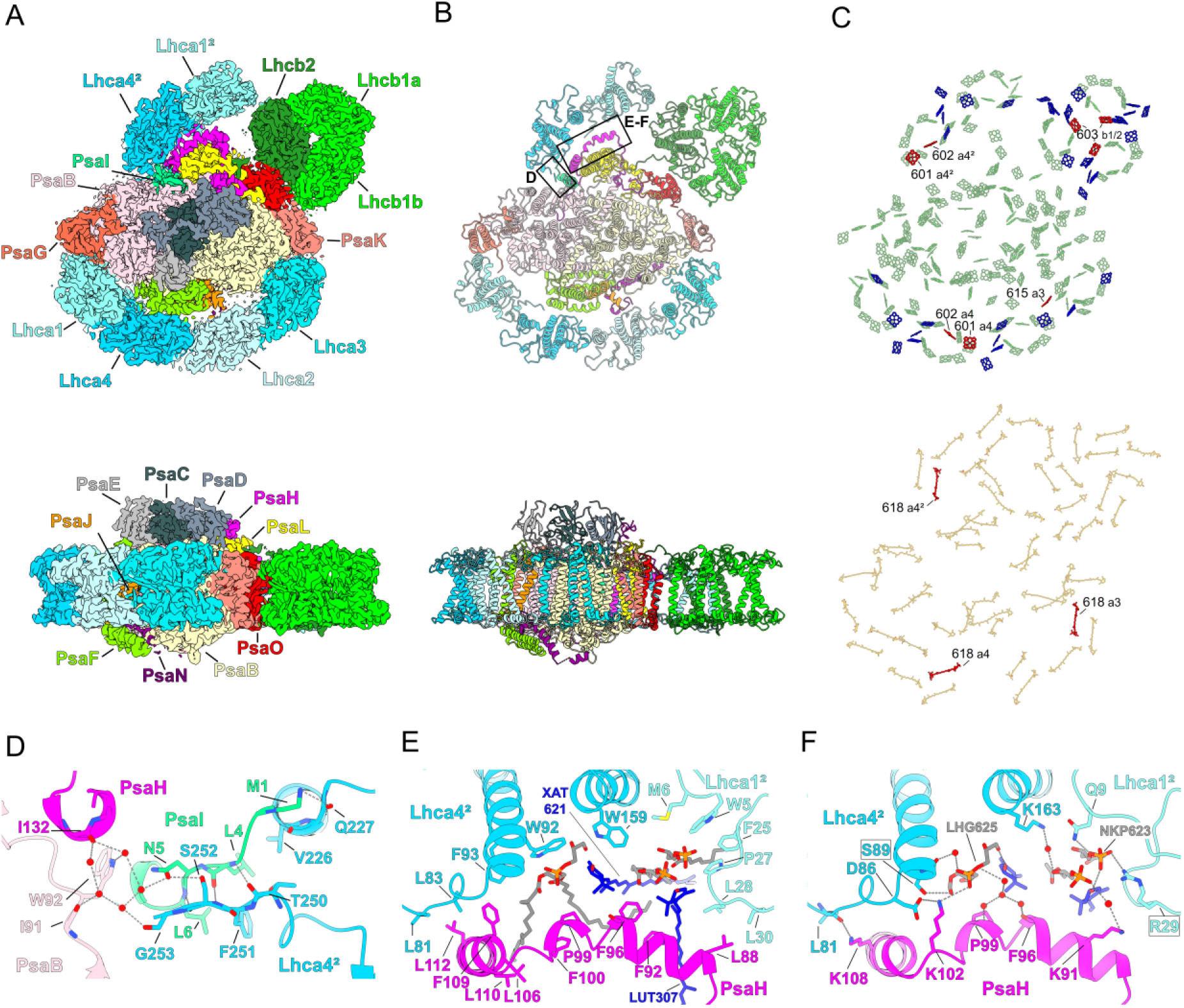
The overall architecture of *P. oceanica* L-PSI-LHCII. **A.** Stromal view (top) and side view (bottom) of the cryo-EM density map of the *P. oceanica* L-PSI-LHCII highlighting the occupancy and identity of each subunit with different colors. **B**. Stromal view (top) and side view (bottom) of the atomic model of the L-PSI-LHCII in ribbon representation. The areas indicated by black squares are enlarged in (D) to (F). **C.** Distribution of chlorophyll molecules in the L-PSI-LHCII (top). Chl *a* is shown in green, conserved Chl *b* in blue, and reassigned Chl *b* in red. For clarity, phytol chains were omitted. Distribution of carotenoid molecules in the L-PSI-LHCII (bottom). Conserved carotenoids are shown in orange and reassigned positions in red. **D-F.** Atomic model of the domain connecting the Lhca1²-Lhca4² with the core in the L-PSI-LHCII. Hydrogen bonds and salt bridges are depicted as dashed lines.

### Attachment of an additional Lhca1-Lhca4 dimer to the core complex

The Lhca1²–Lhca4² dimer attaches to the core via a large interface involving PsaH, Lhca1², and Lhca4², and a smaller one involving Lhca4², PsaH, PsaI, and PsaB (**Figure 1A and 1B**). The small interface involves hydrogen-bonding and Van der Waals contacts that organize the terminal loops of both Lhca4² and PsaI, together with ordered water molecules bridging the PsaI-Lhca4² interaction with PsaH and PsaB (**Figure 1D)**. On the large interface, Van der Waals interactions (**Figure 1E**) as well as salt bridges and hydrogen bonds (**Figure 1F**) stabilize protein–protein contacts between PsaH and Lhca4² whereas this type of interaction mostly links Lhca1² to PsaH through cofactor–protein contacts. Interestingly, Lhca4 establishes a protein–protein contact with PsaL analogous to that formed between Lhca4² and PsaH (**Supplemental Figure 3**), suggesting that Lhca4 already contains the determinants required to bind two distinct positions.

Sequence analysis of the corresponding domains in the small (**Supplemental Figure 4A**) and large interface (**Supplemental Figure 4B**) revealed only two conserved substitutions in seagrasses, affecting the large interface: Asn89→Ser in Lhca4 is predicted to maintain the same interactions through model-based mutagenesis (**Supplemental Figure 5A**), and Gly29 →Arg (Glu in *Z. marina*) in Lhca1 is predicted to only disrupt the hydrogen bond between the side chain and the glycerophospholipid NKP623 (**Figure 1F**). Therefore, the recruitment of the additional Lhca1²–Lhca4² dimer in seagrasses may rely on antenna stoichiometry as previously suggested (Ferroussier et al., 2025), rather than seagrass-specific structural motifs.

### Blue light adaptation in the LHCII

The phosphorylated LHCII trimer occupies the canonical PSI-binding site as in the PSI–LHCII state-transition complex of land plants (Ferroussier et al., 2025; Pan et al., 2018) (**Supplemental Figure 5B**). Furthermore, a Chl substitution previously predicted (Ferroussier et al., 2025) was confirmed at position 603 in all Lhcb monomers of the bound LHCII (**Figure 1C and Figure 2**). This assignment is supported by the seagrass-specific Tyr193, which stabilizes Chl *b*603 through a hydrogen bond with its formyl group, contrasting with other species that have phenylalanine stabilizing the C7 methyl of Chl *a*.

**Figure 2:**
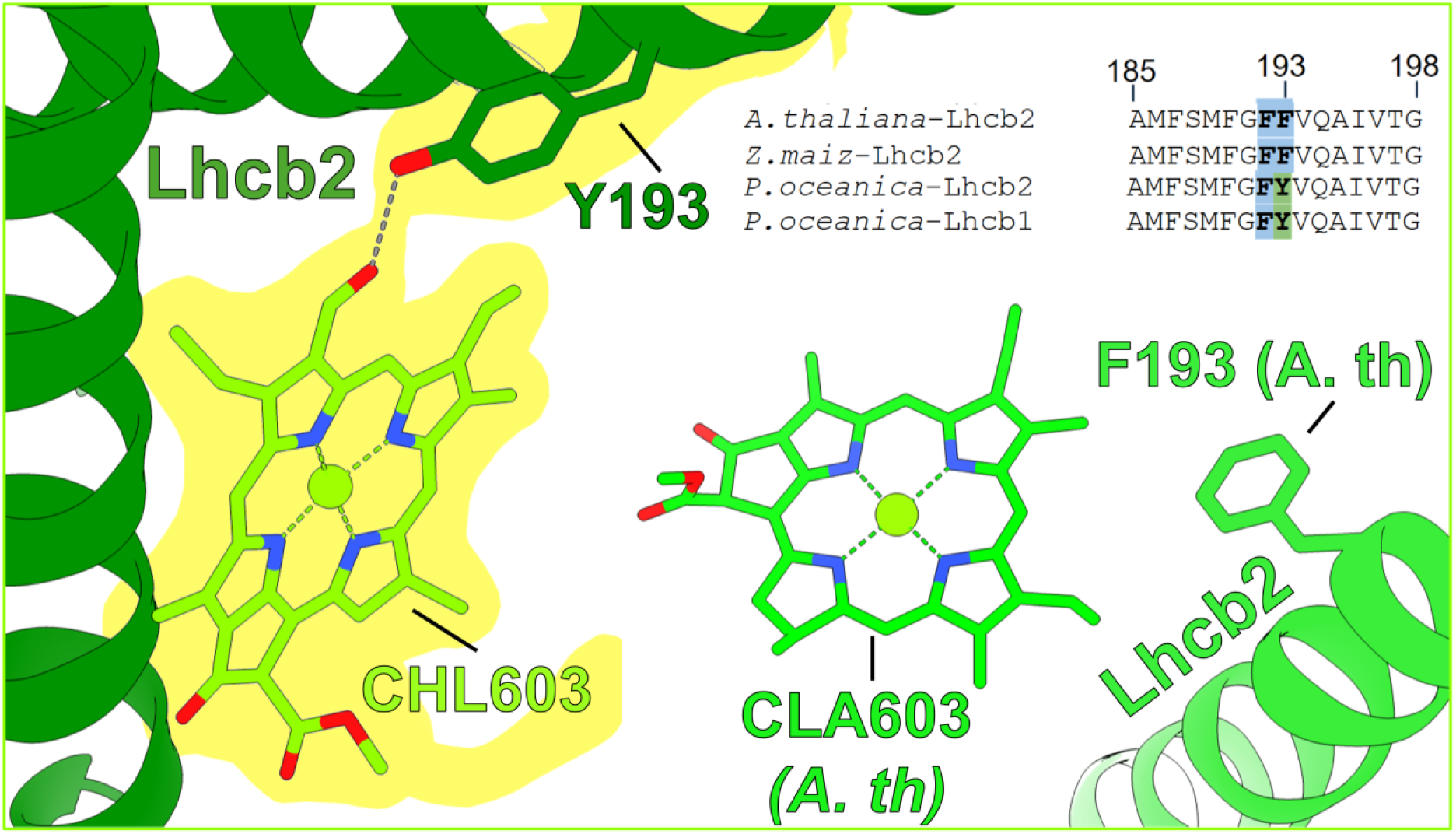
Chl *b* substitution in the bound LHCII. The atomic model with cryo-EM density for the Chl *b* at position 603 is shown interacting with Tyr 193 in Lhcb2 via the formyl group of Chl *b*. The *A. thaliana* PSI model (9GBI) is shown with its equivalent domain, together with the local sequence alignment of Lhcb2/1 from representative species. Position numbers are based on the numbering in the L-PSI-LHCII model.

### Blue light adaptation in the LHCI

Within the Lhcas, three canonical chlorophyll-binding sites were assigned as Chl *b* in *P. oceanica*, although modeled as Chl *a* in reference structures (*A. thaliana (Capaldi et al., 2025)*, *F. albivenis (Li et al., 2024)*, *A. sativa (Naschberger et al., 2024)*). These sites correspond to Chl *b*615 in Lhca3 and Chl *b*601/602 in each Lhca4 copy, all located near the far-red domains (**Figure 3A–3E**). Although up to eight Chl *a*-to-Chl *b* substitutions were previously suggested in *P. oceanica* PSI-LHCI (Ferroussier et al., 2025), only these three could be confidently assigned here. Density features consistent with the Chl *b* formyl group was observed at Lhca3 Chl615 and Lhca4 Chl602 (**Figure 3C and 3D**), while the C7 density at Chl601 overlaps the side chain of Asn76 (**Figure 3E**). Notably, the local protein environment around these chlorophylls is largely conserved between *P. oceanica* and the reference species, and density features potentially compatible with Chl *b* can also be discerned at site 601 in *F. albivenis* and *A. sativa*, and at site 602 in *A. sativa* (**Figure 3C–3E**; **Supplemental Figure 6– 8**), suggesting binding flexibility at these sites. Additionally, water molecules bridging Chl *a*609 and Chl *b*608 were modeled (**Figure 3A and 3B**). Although not reported in prior models, they were found in *F. albivenis* upon map reinspection (**Supplemental Figure 9**).

**Figure 3:**
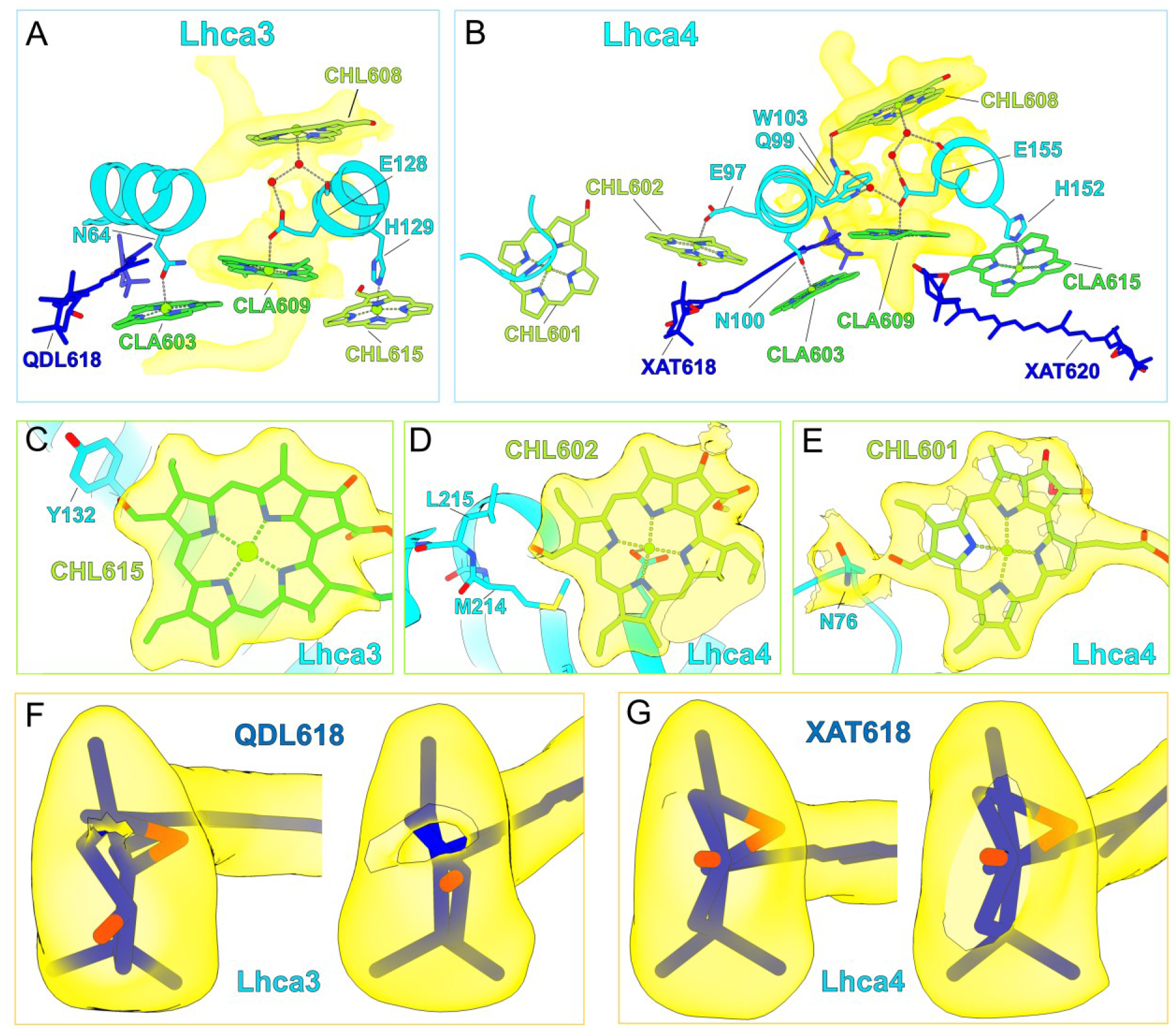
Pigment content in the antenna of PSI. **A-B.** Architecture and co-factors composition near the far-red domains in Lhca3 (A) and Lhca4 (B) from the L-PSI-LHCII. Water molecules are represented by red spheres. Cryo-EM density is shown in yellow only to highlight the residues and the water molecules network bridging Chl *b*608 (CHL618) and Chl *a*609 (CLA609). **C-E** 3D model and cryo-EM density map of the Chl *b*615 (CHL615) site in Lhca3 (C) and Chl *b*601 (CHL601) (D) and Chl *b*602 (CHL602) (E) sites in Lhca4. **F-G.** 3D model and cryo-EM density of the ionone head of carotenoids at position 618 in Lhca3 (F) and Lhca4 (G).

Two carotenoid substitutions were identified at position 618 near the far-red domain in Lhca3 and Lhca4. In Lhca3, density presents the protrusion of one epoxidized ionone ring and the flatness of a non-epoxidized β-ionone ring on the opposite side, consistent with antheraxanthin (**Figure 3F**), its epoxide potentially stabilized by hydrogen bonding with Ser75, unique to *P. oceanica* (**Supplemental Figure 10A**). In Lhca4, density at site 618 displays two epoxide protrusions, consistent with violaxanthin (**Figure 3G**), whose accommodation may be facilitated by reduced steric hindrance caused by the Met→Val substitution at position 111 (**Supplemental Figure 10B**). The local environments and densities suggest lutein at these sites in the other reference species (**Supplemental Figure 10C–10H**), although the available resolution does not allow definitive assignment in reference species. Both antheraxanthin and violaxanthin may contribute to the photoprotective plasticity proposed for *P. oceanica* (Procaccini et al., 2017), while violaxanthin absorption is slightly more blue-shifted compared to lutein (Ashenafi et al., 2023), matching the deep seawater irradiance window. Together with the local protein environment in *P. oceanica*, these observations suggest that both pigments are specific to this species.

### Protein environment of the far-red domains

In *P. oceanica*, the axial ligands of Chl *a*603 and Chl *a*609 in Lhca3 and Lhca4 are Asn and Glu, respectively, as in species exhibiting either a moderate red shift (P*. sativum*, ∼733 nm) or a pronounced red shift (*F. albivenis*, ∼745 nm) (**Figure 3A and 3B**; **Supplemental Figures 11 and Figure 12**). The overall structures of Lhca3 and Lhca4, including their chlorophyll-binding sites, are largely conserved in those species.c(**Figure 4A**). Because the spectral properties of red forms mainly depend on local intrinsic features (Morosinotto et al., 2003; Rankelytė et al., 2024), we compared the molecular environment of the red forms in *P. oceanica* to those of these species.

**Figure 4:**
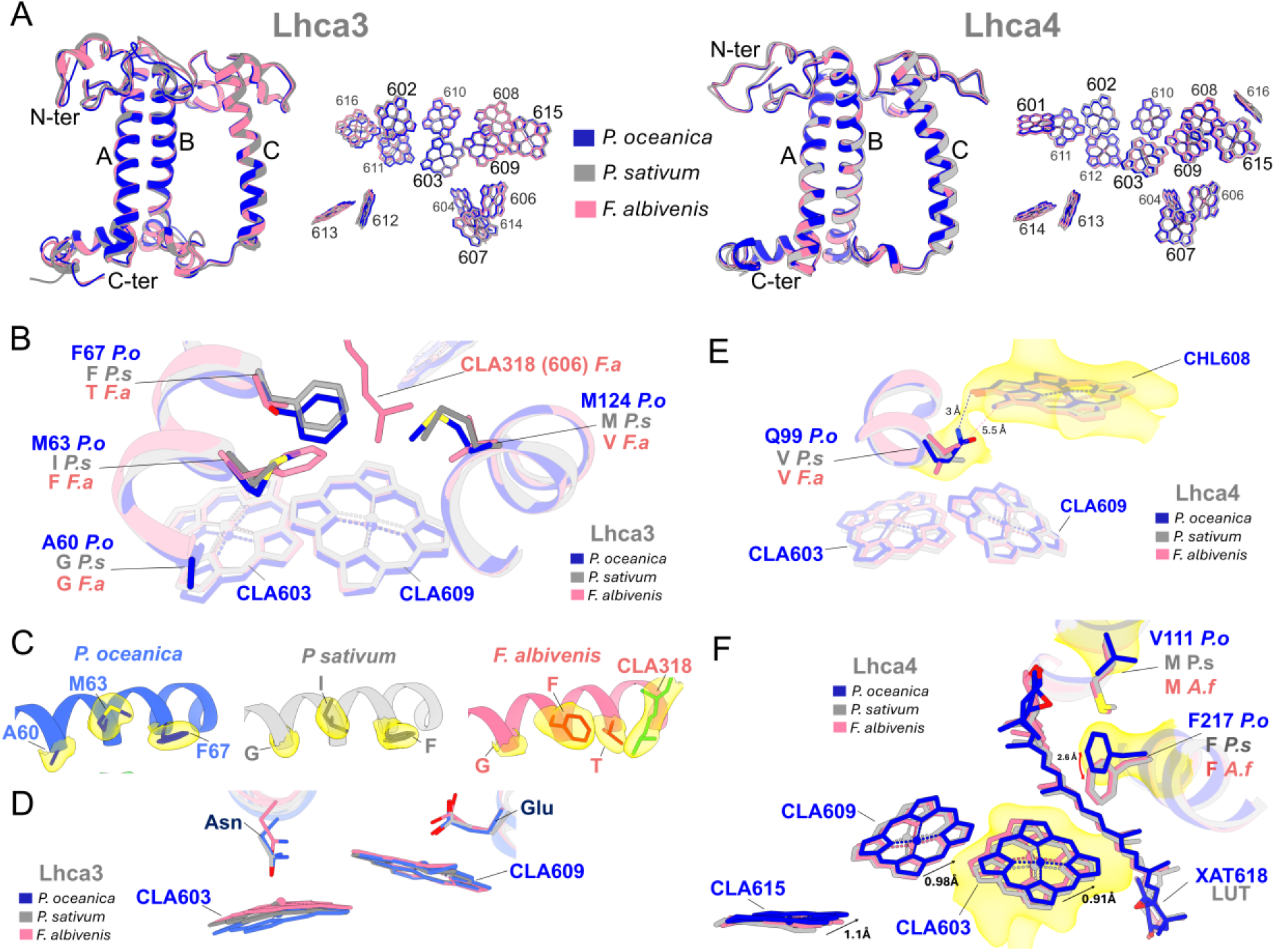
Comparison of the protein environment around the red-forms in Lhca3 and Lhca4. **A**. Superposition of protein structures and chlorophylls arrangement in Lhca3 (left) and Lhca4 (right) from *P. oceanica*, *P. sativum* (PDB:7DKZ) and *F. albivenis* (PDB: 8WGH). The three conserved transmembrane helix are indicated by the letter A, B and C. **B.** Superposition of the atomic model of the L-PSI-LHCII with atomic models from *P. sativum* and *F. albivenis* focused on the red-form domain in Lhca3. Only the non-conserved residues located in the direct environment of the red-forms are highlighted. CLA318 is named as determined by authors with the equivalent position number in *P. oceanica* into brackets **C**. Atomic model and cryo-EM density (yellow) comparison at positions 60, 63 and 67 in *P. oceanica* and equivalent positions in *P. sativum* and *F. albivenis* **D.** Superposition of the Chl *a*603 (CLA603) and Chl *a*609 (CLA609) domain in different species. The 3D models were aligned on Chl *a*609 from *P. oceanica*. **E.** Superposition of the atomic model of the L-PSI-LHCII with atomic models from *P. sativum* (PDB:7DKZ) and *F. albivenis* (PDB: 8WGH) focused on the red-form domain in Lhca4. Only the non-conserved residues located in the direct environment of the red-forms are highlighted. **F**. Same as D but extended to the pigment and residues bordering the far-red domain in Lhca4. Shift lengths are given as the distance between residues in *P. oceanica* and the most distant corresponding superposed residues in the other species.

In Lhca3, *P. oceanica* differs from *P. sativum* by Ala60 and Met63 (**Figure 4B**). Compared with Gly60 and Ile63 in *P. sativum*, Ala60 likely increases steric pressure on Chl *a*603 through its methyl side chain, whereas Met63 introduces an unbranched thioether side chain which may relax constraints above Chl *a*609 (**Figure 4B and 4C**; **Supplemental Figure 13**). These substitutions coincide with a downward tilt of the Chl *a*603 tetrapyrrole ring relative to Chl *a*609, resulting in a less parallel arrangement and greater separation (**Figure 4D**). In Lhca4, Gln99 lies above Chl *a*609 and forms a hydrogen bond with the C13 ketone of Chl *b*608, an interaction absent from other species carrying Val at this position (**Figure 4E**; **Supplemental Figure 14**). The co-facial, near-parallel arrangement of Chl *a*608 and Chl *a*609 together with the structured chlorophyll–ligand–water network linking them (**Figure 3A and 3B**), suggests that changes at this site could influence the excitonic environment of Chl *a*609. A second feature is the Met→Val substitution at position 111, which reduces local steric constraints and allows Phe217 to adopt an alternative rotamer. These changes are accompanied by displacement of Chl *a*603, Chl *a*609, and Chl *a*615 by up to ∼1 Å toward violaxanthin 618 (**Figure 4F)**.

Remarkably, *F. albivenis* differs from *P. sativum* and *P. oceanica* at positions 63 (Phe), 67 (Thr), and 124 (Val) in Lhca3 (**Figure 4B and 4C**). The bulky Phe63 overlaps Chl *a*609, whereas the smaller Thr67/Val124 residues create space for the phytol chain of Chl *a*318 (606 in *P. oceanica*) to insert above Chl *a*609 (**Figure 4C**; **Supplemental Figure 15A**). Such feature was absent from the original model and was also visible upon reinspection of the *A. sativa* density map (PDB: 8BCV; **Supplemental Figure 15B**). These changes are accompanied by an upward shift of the α-helix hosting the axial ligand of Chl *a*603 in *F. albivenis*, together with tilting of its macrocycle and reduced Chl *a*603–Chl *a*609 separation relative to *P. sativum* and *P. oceanica* (**Fig. 4D**).

Excitonic coupling (EC) alone does not account for the red forms, but was used as a quantitative descriptor of structural changes affecting pigment-pigment interactions. Thus, the observed rearrangements are associated with a reduction of the |Chl 603–Chl 609| excitonic coupling (EC) by approximately 12% in Lhca3 and 18% in Lhca4 in *P. oceanica* compared to *F. albivenis*. Conversely, the |(Chl *a* 603-Car 618)| EC is higher by 21% in Lhca3 and 13% in Lhca4 in *P. oceanica* compared to *F. albivenis* (**Supplemental Table 2)**.

### Diversification of the far-red domains

To identify the amino acid underlying PSI spectral diversity, we next examined Lhca3 and Lhca4 protein sequences in species spanning the major higher-plant lineages. Species were grouped by far-red emission into three classes centered on *P. oceanica* (∼720 nm), *P. sativum* (∼734 nm), and *F. albivenis* (>740 nm) plus one class represented by the lycophytes *S. denticulata* (726 nm (Capaldi et al., 2025)). This analysis again highlighted the four sites in Lhca3 and the two sites in Lhca4 described in **Figure 4**, whose residue identity or similarity delineates each far-red emission class (**Figure 5**; **Supplemental Figure 11** and **Figure 12** for the complete sequence analysis).

**Figure 5:**
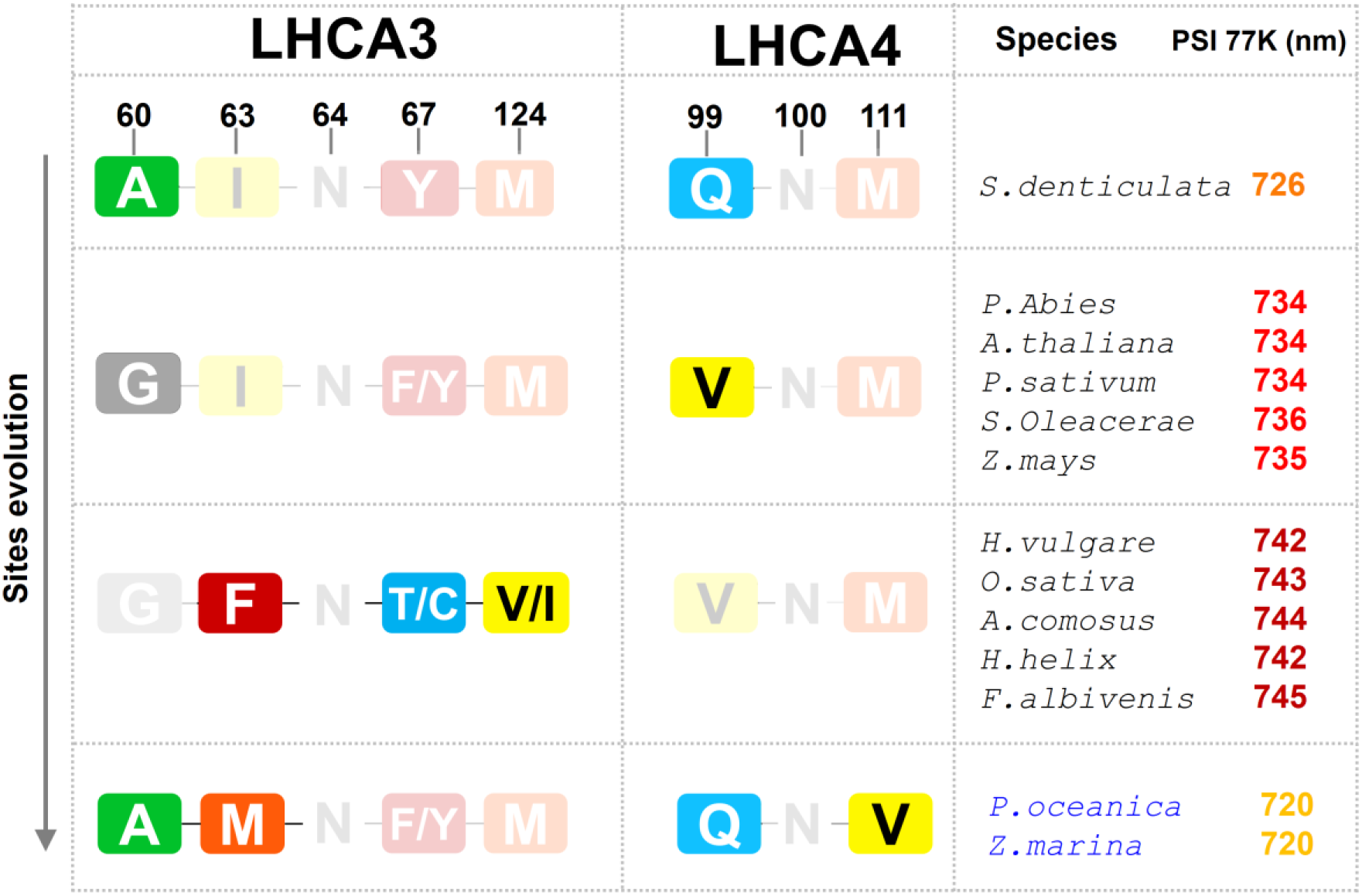
Evolution of the far-red domains in Lhca3 and Lhca4. The far-red domains of Lhca3 and Lhca4 from representative higher plant species were sorted and grouped based on the conservation of the residues identified in Figure 4. The corresponding PSI 77 K fluorescence emission values are shown alongside. Position numbers are based on the numbering in the L-PSI-LHCII model.

More specifically, Gly60 in Lhca3 and Val99 in Lhca4 recur in all land spermatophytes (seed plants), including both red-(∼734 nm) and strongly red-shifted (>740 nm) species. By contrast, Ala60 and Gln99 are conserved in the mildly red-shifted lycophyte *S. denticulata* (726 nm) and in the two seagrasses (720 nm), indicating reversions toward an ancestral-like state in *P. oceanica* and *Z. marina*. These seagrasses are further distinguished by Met63 in Lhca3 and Val111 in Lhca4. Together, this pattern suggests that the Ala→Gly and Gln→Val substitutions may have substantially contributed to the far-red shift during the evolution of land plants. Finally, Phe63, Thr67 (Cys in *H. helix*), and Ile124 are conserved in all strongly red-shifted species, defining the most divergent state relative to less red and bluer species, and consistent with the distinctive structural features observed in *F. albivenis* compared with *P. sativum* and *P. oceanica* (**Figure 4**). Interestingly, selected land and freshwater Alismatales retain the same far-red emission and motifs as red-shifted land species (∼734 nm) in both Lhca3 and Lhca4 (**Supplemental Figure 16A**). By contrast, the seagrass *H. stipulacea*, despite emitting at 719 nm (Sharon et al., 2011), retains the land-plant-like Ile63 in Lhca3 but carries Cys at position 60, predicted to exert stronger steric pressure on Chl *a*603 than Ala and potentially compensate for the absence of Met63 (**Supplemental Figure 16B and 16C**). Together, these observations indicate that the diversity of far-red emission in higher plants maps onto variation in the composition of the same recurrent amino acid sites in Lhca3 and Lhca4.

## DISCUSSION

One of the fundamental questions regarding the evolution of photosynthesis is how spectral diversification can be achieved by a highly conserved photosynthetic complex, while preserving both its photochemical efficiency and its structural integrity. As revealed by our 1.9 Å cryo-EM structure of the L-PSI-LHCII supercomplex in *P. oceanica*, the adaptation of higher plants PSI relies on localized remodeling of a limited number of sites allowing for spectral tuning, rather than on a profound reorganization of the photosystem’s architecture. Indeed, the remarkable spectral specialization of *P. oceanica*, characterized by increased capture of blue and cyan light as well as a significant attenuation of far-red light absorption, does not substantially alter either the PSI core or the overall structure of the LHCI antenna.

The recruitment of the additional Lhca1-Lhca4 dimer in the L-PSI-LHCII supercomplex occurs through interaction surfaces that are highly similar to those described in terrestrial angiosperms. Both the PSI–LHCII and PSI–Lhca1/Lhca4 interfaces are mediated by residues that are highly conserved across seagrasses and land angiosperms (Pan et al., 2018; Wu et al., 2023), suggesting that the structural capacity to assemble the L-PSI-LHCII supercomplex was already present within the angiosperm PSI architecture and did not require the evolution of novel structural features in seagrasses (**Figure 1**). Instead of major structural modifications, adaptation appears to involve changes in the relative abundance of pre-existing photosynthetic components including: (i) a higher PSI content, (ii) enhanced accumulation of Lhca1/Lhca4 and (iii) high LHCII content, a set of traits so far observed only in seagrasses expressing the L-PSI-LHCII (Ferroussier et al., 2025). The selective recruitment of this complex in seagrasses therefore appears to reflect differences in gene expression rather than the emergence of lineage-specific structural innovations. These findings support the view that seagrass adaptation to the seawater light environment primarily involved the differential recruitment of a conserved ancestral photosynthetic “toolkit”, with evolutionary change arising from modulation of existing components rather than the set up of new structural features.

Although enhanced blue–cyan light harvesting is associated with Chl *a*-to-Chl *b* substitutions in both LHCI and LHCII (Ferroussier et al., 2025), the Chl *b* assignments in the L-PSI-LHCII revealed distinct pigment substitution mechanisms in the two antenna systems (**Figure 2 and 3**). The reassigned Chl sites in the LHCI occurred within a protein environment largely conserved across angiosperms, suggesting that these sites can accommodate either Chl *a* or Chl *b* in seagrasses, and more broadly in angiosperm LHCI. Such flexibility could allow pigment composition to vary with Chl *a/b* availability, as suggested in mosses (Gorski et al., 2022). This could facilitate acclimation to contrasting light environment in land habitats and along the depth gradient or during seasonal cycles in seagrasses (Ferroussier et al., 2025). In contrast, Chl *b* incorporation in LHCII appears constrained by seagrass-conserved amino-acid substitutions, notably Tyr193, which stabilizes Chl *b* at site 603 in each Lhcb1/2 monomer. This site lies in a photoprotective domain in land plant, where Chls *a*602/*a*603 contribute to excess-energy quenching through adjacent lutein (Agostini et al., 2021; Carbonera et al., 2022), that is suppressed when Chl *b* occupies the site in recombinant land-plant LHCII (Kleima et al., 1999). Thus, Chl *b*603 incorporation in seagrasses may involve a trade-off between reduced photoprotection and a constitutively enhanced blue–cyan light harvesting in the mobile LHCII, complementing the likely-tunable Chl *b* enrichment in the LHCI. Interestingly, Chl replacement in the LHCII also occurred in depth-adapted green algae, such as *Bryopsis corticulans* (Li et al., 2023) and *Codium fragile* (Seki et al., 2022), which can incorporate Chl *b* at site 602. This further indicates that Chl *b* incorporation around sites 602/603 may represent a recurrent trait of seawater adaptation in green plants.

The strong attenuation of the red forms in L-PSI-LHCII further enhanced absorption of higher energy photons and provided the energetic landscape compatible with both a large antenna size and efficient energy trapping (Ferroussier et al., 2025). A similar combination occurred in green algae, where blue-shifted red forms compensate for PSI antenna enlargement (Qin et al., 2015; Le Quiniou et al., 2015). In *P. oceanica*, however, PSI retained the same LHCI architecture as higher plants, composed of the four canonical Lhca’s paralogue (Lhca1-4), and red forms were attenuated despite retaining the Chl *a*603–Chl *a*609 pair and the conserved His→Asn substitution in Lhca3/4. Since restoring His in *A. thaliana* weakens Chl *a*603–Car618 coupling (Capaldi et al., 2025), red forms may have been attenuated in seagrasses while preserving land-plant-like carotenoid-mediated photoprotection (Carbonera et al., 2005; Alster et al., 2024), particularly essential for shallow meadows exposed to high light and caustic focusing (Hieronymi and Macke, 2012; Procaccini et al., 2017). Thus, rather than reverting PSI towards the ancestral His state, seagrasses attenuated red forms through local remodeling of pigment–protein interactions within the canonical Chl-ligand system. A major finding of this study is the identification of recurrent amino-acid sites surrounding the far-red domains of Lhca3 and Lhca4 that differ systematically between blue-shifted and red-shifted species (**Figure 4 and 5**). Variations at these positions are associated with distinct local structural environments and subtle rearrangements of the red forms geometry (**Figure 4**). Across the species analyzed, similar fluorescence spectral ranges systematically correlate with comparable residue states or structurally convergent substitutions (**Figure 5**, **Supplemental Figure 16)**. Altogether, these data support a common structural basis underlying PSI spectral diversification in higher plants. The identified sites therefore emerge as candidate evolutionary hotspots for red-form tuning, and may provide targets for future red-form engineering in crops.

Comparison of the far-red domains in Lhca3 and Lhca4 revealed different evolutionary trajectories in the two paralogues. Red forms in Lhca3 traces back to mosses where the co-axial ligand His was substituted by Asn (Alboresi et al., 2008). This event was followed by the diversification of the protein environment around the red-forms, notably in seed plants, where an Ala-to-Gly substitution at position 60 correlates with emission shifted to ∼735 nm, while three additional substitutions in shade-adapted dicots and most monocots associates with emission beyond 740 nm (**Figure 4 and 5**). By contrast, the far-red domain of Lhca4 remains comparatively constrained, showing fewer variable sites across lineages and a conserved composition in red-shifted and strongly red-shifted species. This pattern suggests that, although Lhca4 is the dominant red-emitting paralogue in *A. thaliana* (Wientjes et al., 2012), Lhca3 became the main contributor to far-red emission in strongly red-shifted species. Moreover, the remodeling in Lhca3 extended beyond the protein environment of the red forms, with insertion of the Chl *a*606 phytol chain above Chl *a*609 in *F. albivenis,* adding a non-protein structural element to the far-red domain. Together, these observations identify Lhca3 as a major driver of PSI far-red diversification, highlighting how structural flexibility among LHCI paralogues shaped PSI spectral adaptation during plant evolution.

In conclusion, the L-PSI-LHCII supercomplex illustrates how evolutionary innovation can arise within a highly conserved photosynthetic framework. By combining canonical PSI architecture with a limited set of seagrass-specific modifications, *P. oceanica* achieves extensive spectral reprogramming without compromising photochemical performance. The findings presented here establish PSI as a structurally conserved yet spectrally flexible complex, able to adapt light harvesting to contrasting photic environments through parsimonious and localized architecture remodelling. Such evolutionary plasticity enables higher-plant PSI to adapt to a wide variety of spectral environments while preserving photochemical integrity and efficient energy trapping.

## METHODS

### Sample collection

*Posidonia* oceanica shoots (sheath bundles, and leaves) were harvested at 26m depth in the south of Frioul Island, Marseille, France (Coordinates 43°16’11” N 5°17’32” E). The samplings were carried out within the framework of a protected species exemption after obtaining a prefectural decree authorizing these samplings. Immediately after harvesting, plants were kept in the dark in cold seawater. Leaves were separated from bundles at 4°C under dim green light, and epiphyte organisms were removed by washing with seawater.

### Isolation and separation of photosystem supercomplexes

Thylakoid membranes were prepared as previously described (Charras et al., 2024). The chlorophyll *a/b* ratio and chlorophyll content of the thylakoid membranes were measured as described in (Chazaux et al., 2022).

### L-PSI-LHCII purification

Purified thylakoids (2 mg Chl/ml) were solubilized with a final concentration of 4% (w/v) **α**-DDM for 20 min in a buffer containing 40mM Hepes pH 7.4, then centrifuged at 21,000 g. The supernatant was loaded onto a sucrose gradient (0.75 M sucrose, 40 mM Hepes pH 7.4, 0.015% **α**-DDM), formed by freeze-thawing. After ultracentrifugation, the target band was collected, dialyzed against 40 mM Hepes pH 7.4 and concentrated. The fraction underwent a second step of sucrose gradient ultracentrifugation to partially remove PSI-LHCI contaminants. The target band was collected followed by dialysis and concentration.

### Protein sequence analysis

Lhca3 and Lhca4 protein-coding sequences from the representative species analysed in this study were retrieved using NCBI tBLASTx searches with *A. thaliana* Lhca3 and Lhca4 as queries. Candidate sequences were verified by inspection of conserved Lhca motifs. Multiple sequence alignments were performed using T-Coffee (Notredame et al., 2000).

### Cryo-EM data collection and processing

The L-PSI-LHCII sample with a chlorophyll concentration of 2.5 mg ml^−1^ was applied to glow-discharged (GloQube Quorum, 30 sec, 35mA) holey carbon grids (Quantifoil 300 Au R1.2/R1.3, Electron Microscopy Sciences). For vitrification a Vitrobot Mark IV (Thermo Fisher Scientific) was used at 4°C and 100% humidity with a chosen blotting time of 2.5 sec. The data was collected using a 300 kV Titan Krios G4 Microscope (Thermo Fisher Scientific) equipped with a Falcon 4i direct electron detector (Thermo Fisher Scientific) and a Selectris Energy filter (Slit width 10 eV) energy filter. 23,818 movies were recorded at a nominal magnification of 165,000x, corresponding to a calibrated pixel size of 0.729 Å and a total dose of 40 e Å^-2^ per movie. The defocus range was set from −0.5 to −2.1 µm in 0.2 µm steps. The data statistics are shown in **Supplemental Table 1**.

Processing was performed using cryoSPARC 4.7.1 (Punjani et al., 2017). Blob and template pickers were used to pick a subset of particles (box size of 400 px, binned to 100 px) which were extracted and subjected to 2D classification and ab initio reconstruction using 24,161 particles. The low-resolution map was used as an input reference for subsequent rounds of multi-class heterogeneous refinement together with several junk classes (in total seven heterogeneous refinement jobs). In total, 6,720,319 particles were picked and used as input for this series of heterogeneous refinement steps. Using this procedure, we enriched a subset of 286,004 high-quality particles, which were extracted unbinned with a box size of 600 px. Non-uniform (NU) refinement followed by global and local CTF refinement and an additional NU refinement yielded a reconstruction of the L-PSI–LHCII structure at 2.1 Å resolution. Further 3D classification, followed by reference-based motion correction in cryoSPARC and a final NU refinement, resulted in a map with a resolution of 2.0 Å based on 221,716 particles.

The particle information was transferred into a Star file format (UCSF pyem v0.5. Zenodo) and imported into RELION 5.0.0b (Scheres, 2012; Kimanius et al., 2021). The movies in RELION were motion corrected and dose weighted using MotionCor2 (Zheng et al., 2017). The parameters for the Contrast Transfer Function (CTF) of each particle were used as estimated by cryoSPARC. Particles were re-extracted and processed in RELION using a box size of 450 pixels and a pixel size of 0.729 Å.

3D Refinement was performed followed by Post processing and CTF refinement (Rohou and Grigorieff, 2015; Zhang, 2016). Another round of 3D refinement and post processing was performed followed by Bayesian polishing (Scheres, 2012), 3D refinement, post processing and a second round of CTF refinement. After an additional round of post-processing, Bayesian polishing, and 3D refinement, a 3D classification with three classes was performed. Two classes (1 and 3), which showed higher map quality, were selected and subjected to 3D refinement followed by post-processing to obtain the final consensus map at 1.92 Å resolution from 182,928 particles.

To improve the resolution of the PSI core, LHCII, and the extra Lhca1–Lhca4 complex, we first performed masked refinement followed by signal subtraction outside the respective masks for all three map regions. Subsequently, 3D refinement was carried out for each region, followed by post-processing to obtain improved resolutions for the three map regions (PSI core: 1.87 Å; LHCII: 2.17 Å; Lhca1–Lhca4 dimer: 2.21 Å).

### Model building and refinement

The structure of PSI-LHCII of maize (PDB ID: 5ZJI) was a rigid body fitted into the 1.9 Å map using Chimera v 1.17 (Pettersen et al., 2004). Then, SWISS-Model (Waterhouse et al., 2018) and Alpha Fold 2 (Jumper et al., 2021) were used to generate models of each protein in the complex. The models were superposed chain by chain onto the maize template structure using Coot v9.8.98 (Emsley et al., 2010). All protein residues as well as pigments were manually real space refined using *Coot* with locally optimized map weights. The exchange of Chl *a* to a Chl *b* was only modelled when it was supported by the density map and by a favorable chemical environment of the surrounding. New pigments were identified by placing possible candidates into the density. The pigments which matched the density and chemical environment best were then selected and built. All water molecules were placed manually using *Coot*. For all ligands in the model, restraint files were generated using the Grade server (http://grade.globalphasing.org). If required, restraint files were adopted manually.

The model was refined using Real-Space-Refine from the PHENIX suite (Afonine et al., 2018). For the refinement, restraint files generated as described above were used. Distance .edit file for chlorophyll coordinating residues were generated by Ready-set in PHENIX and used throughout the refinement. Hydrogen atoms were added to the model prior to refinement using Ready-set. Various weight parameters were tried out and optimized. Multiple rounds of validation and real-space refinement were carried out using MolProbity and *Coot*. The structure was validated using the PDB Validation server (https://validate-rcsb-2.wwpdb.org). All figures were prepared using ChimeraX (Pettersen et al., 2021).

### Energy transfer and excitonic coupling computation

Excitonic couplings (ECs) were estimated using a point-dipole approximation (PDA) following Capaldi *et al*. For chlorophylls, point dipoles were positioned at the geometric center of the four nitrogen atoms of the chlorin ring (NA, NB, NC and ND). Chl Q_y_ and Q_x_ transition dipoles were defined along the NB–ND and NA–NC axes, respectively. For carotenoids, dipoles were positioned at the center of the conjugated system and oriented along the polyene chain axis. In our models, this corresponded to atom C15 for lutein and violaxanthin, and to atom C23 for antheraxanthin, with dipole directions defined along C12–C33 for lutein/violaxanthin and along C18–C26 for antheraxanthin.

ECs were calculated as 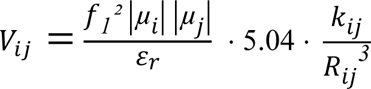

where *f_1_* is the local field correction factor, |*µ_i_*| and |*µ_j_*| are the transition dipole moments of pigments *i* and *j*, *ε_r_* is the relative dielectric constant, *R_ij_* is the distance between dipole centers, and *k_ij_* is the orientation factor.

The latter was calculated as: 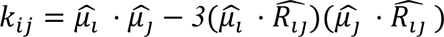

As in Capaldi *et al*., a relative dielectric constant *ε_r_*=2.4 was used, and transition dipole moments were set to 4 D for Chl *a*, 3.4 D for Chl *b*, and 4.5 D for carotenoids, with 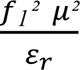 equals 17.6 D² for Chl *a*. Chl–Chl couplings were computed using Q_y_ dipoles, whereas Chl– carotenoid couplings were computed using the Chl Q_x_ dipole and the carotenoid S_2_ dipole. Only absolute EC values were considered and used as geometric descriptors for comparative analysis across species.

### Low temperature fluorescence data

All data were obtained from literature: *P. oceanica* and *Z. marina*: (Ferroussier et al., 2025)*. H. stipulacea*: (Sharon et al., 2011)*. P. abies: (Stolárik et al., 2018). S. denticulata, A. thaliana, A. comosus, O. sativa, Z. mais, S. oleracea*: (Capaldi et al., 2025), (Bos et al., 2023)*. P. sativum*: (Velitchkova and Popova, 2005; Ruban and Johnson, 2009)*. H. helix*: (Zer et al., 2025)*, F. albivenis*: (Li et al., 2024)*. H. vulgare*: (Knoetzel et al., 1998)*, E. nuttallii*: (Maharramov et al., 2015)*, L. minor*: (Liebers et al., 2023).

## ACKNOWLEDGMENTS

We are grateful to Dorian Guillemain, Sandrine Ruitton, Pascal Mirleau, Deny Malengros, Fabrice Garcia, Michel Lafont, Sandrine Chenesseau, Frédérique Legendre, and Bruno Belloni from the OSU Institut Pytheas (Aix Marseille University, CNRS, IRD, IRSTEA, OSU Institut Pythéas, Marseille, France) for the *Posidonia oceanica* sample collection. We are grateful to EMBO for awarding Q.C.F. a Scientific Exchange Grant (number 9348) and we thank Alexey Amunts. We express our deep appreciation to Laurie Casalot, Lea Sylvi, Manon Bartoli, Corinne Valette, and Sophie Guasco for providing essential equipment for sample preparations. We are deeply grateful to Paul Hudson for providing a scientifically enriching research environment.

## Declaration of generative AI and AI-assisted technologies in the writing process

During the preparation of this work, the authors used ChatGPT to improve language and readability. After using this tool, the authors reviewed and edited the content as needed and take full responsibility for the final version of the manuscript.

## FUNDING

This research was supported by the Swedish Foundation for Strategic Research (SSF; grant number ARC19-0051).

## AUTHORS CONTRIBUTIONS

C.J. initiated and developed the research project. Q.C.F., C.J. and A.N. planned and designed the experiments. Q.C.F. collected the plants and performed protein purification. T.A. and AA-A. prepared the sample for Cryo-EM analysis. T.A. and A.A-A. collected the cryo-EM data and processed the data with contribution from A.N.. T.A. carried out BLAST searches and protein modeling. Q.C.F. built the atomic model with contributions of A.N. and T.A.. Q.C.F. and M.S. refined the atomic model and deposited the data. Q.C.F. analysed the structure. Q.C.F. carried out BLAST searches and performed sequence analysis and interpreted the data. Q.C.F. interpreted the structural data. Q.C.F. and C.J. analyzed and interpreted the data with contributions from M.S. and A.N. Q.C.F. prepared the figures. C.J., A.N. and Q.C.F. revised the figures. Q.C.F. and C.J. wrote the manuscript with contribution from A.N., T.A., A.A-A., M.S. and E.H. edited and revised the manuscript.

## COMPETING INTERESTES

Authors declare that they have no competing interests

## DATA, CODE AND MATERIALS AVAILABILITY

The materials generated during the current study are available from the corresponding author on reasonable request. All data necessary to evaluate and reproduce the results in the paper are present in the paper and/or the Supplementary Materials.

## Notes

### Competing Interest Statement

The authors have declared no competing interest.

### Summary of Updates

Improvment prior submittion. The new version incorporates modification in the abstract, Introduction and results sections.

