## Supplementary material for "A new twist on an old idea: The *Posidonia oceanica* Large PSI-LHCI-LHCII supercomplex provides the structural basis of photosystem I spectral diversification in higher plants": Suplemental Figures

### Suplementary Materials

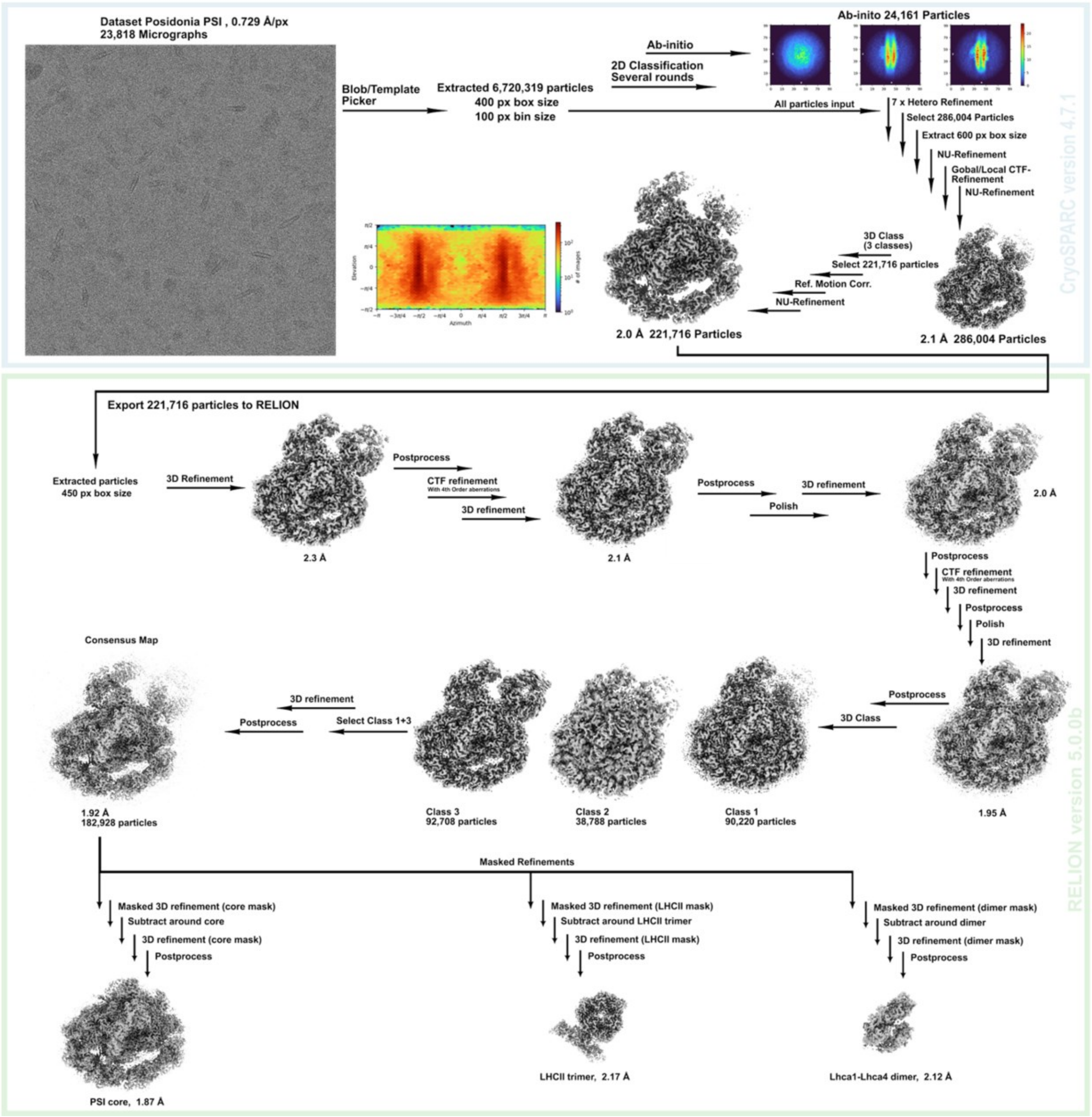

**Supplemental Figure 1. Single-particle cryo-EM data processing workflow used to determine the L-PSI–LHCII structure.**

A representative micrograph is shown. Processing steps performed in cryoSPARC are shown within the light blue box, whereas steps carried out in RELION are shown within the green box. The different job types are indicated next to the arrows. The final map from cryoSPARC before export is shown together with its angular distribution. In RELION, the consensus map was split into three branches for focused refinement of the PSI core region, the LHCII trimer region, and the additional Lhca1-Lhca4 dimer region, each with signal subtraction. Maps are labeled with the particle number and the resulting resolution.

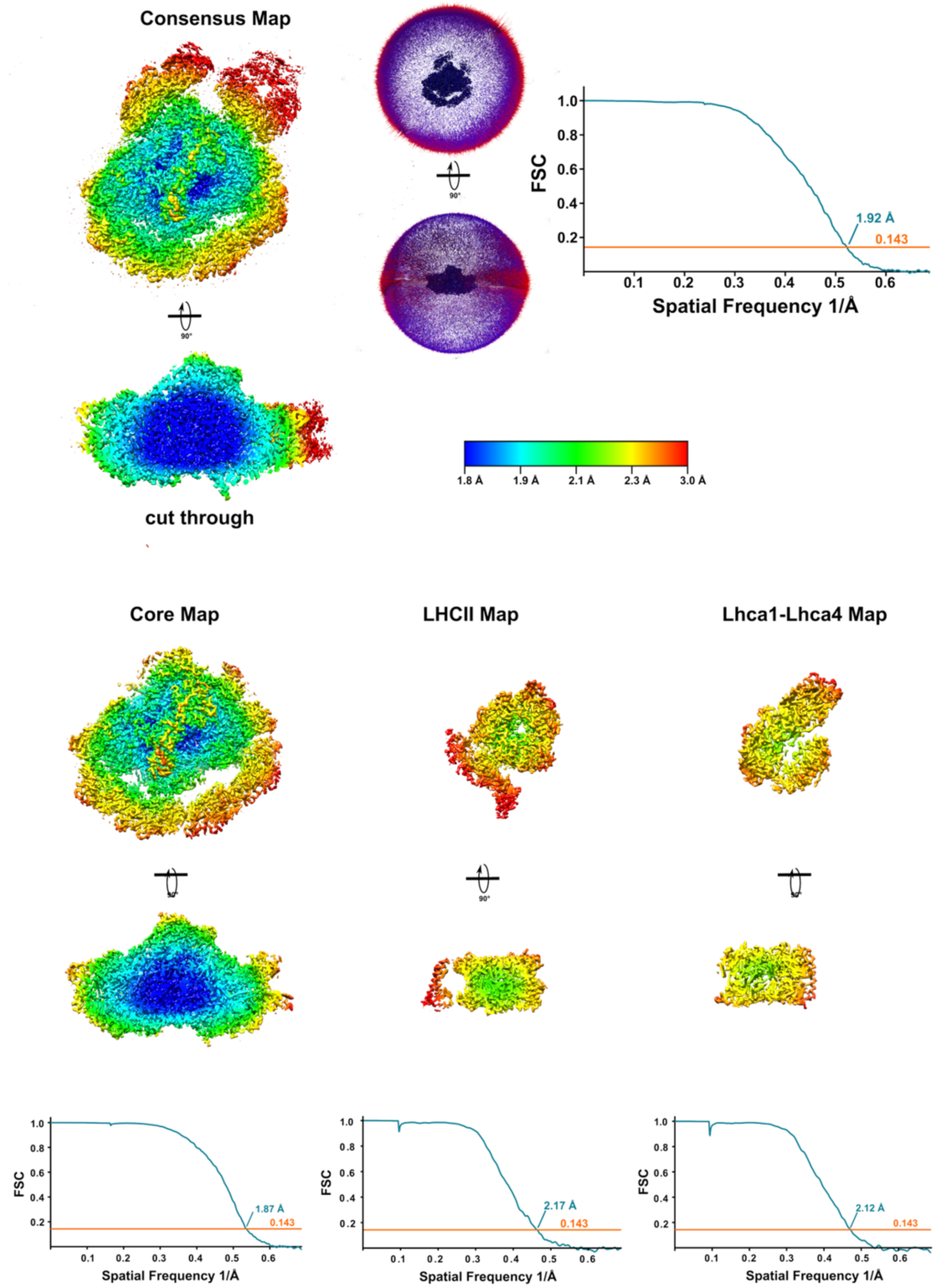

**Supplemental Figure 2. Local resolution, Fourier shell correlations (FSC), and angular distribution of the L-PSI-LHCII structure.**

The consensus map, PSI core map, LHCII map, and Lhca1-Lhca4 map are shown in two orientations in isosurface representation colored by local resolution. The angular distribution is shown for the consensus map. FSC curves for all four maps are provided, with the 0.143 gold-standard cutoff indicated.

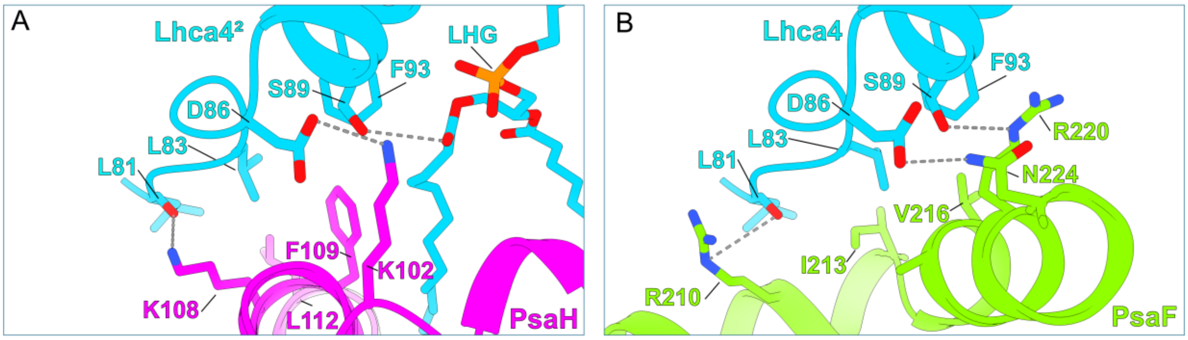

**Supplemental Figure 3.** Comparison of the residues from each Lhca4 copy engaged in the interface with PsaH (**A**) and PsaF (**B**).

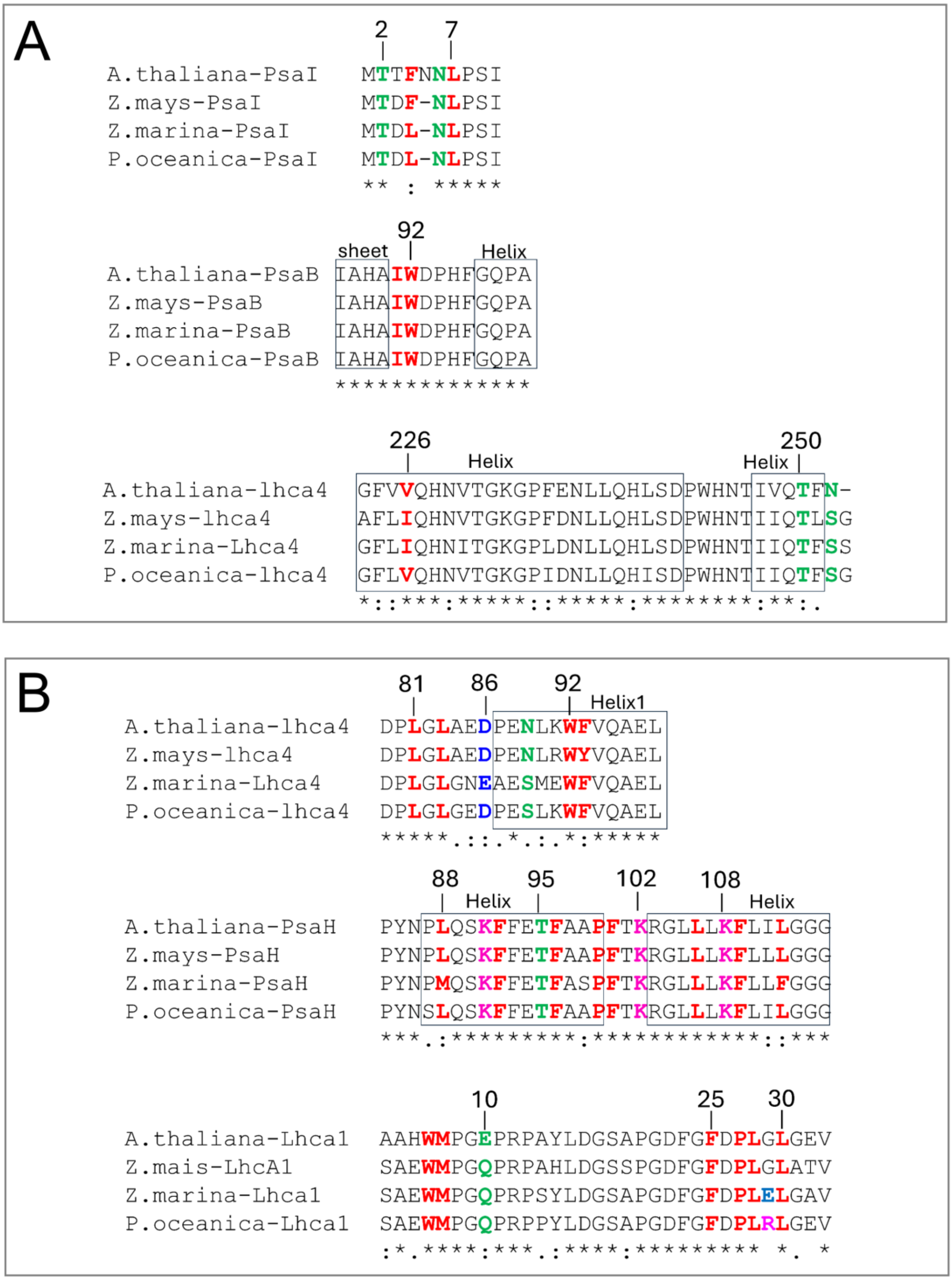

**Supplemental Figure 4. Conservation of the structural determinant of Lhca1²-Lhca4² binding to PSI-core.**

Local protein-sequence alignments of Lhca4, Lhca1, PsaI and PsaH, with residues involved in the (Lhca1²–Lhca4²)–PSI core interface highlighted. The domains involved in the small interface are shown is panel A and the residues from the larger interface are shown in panel B.

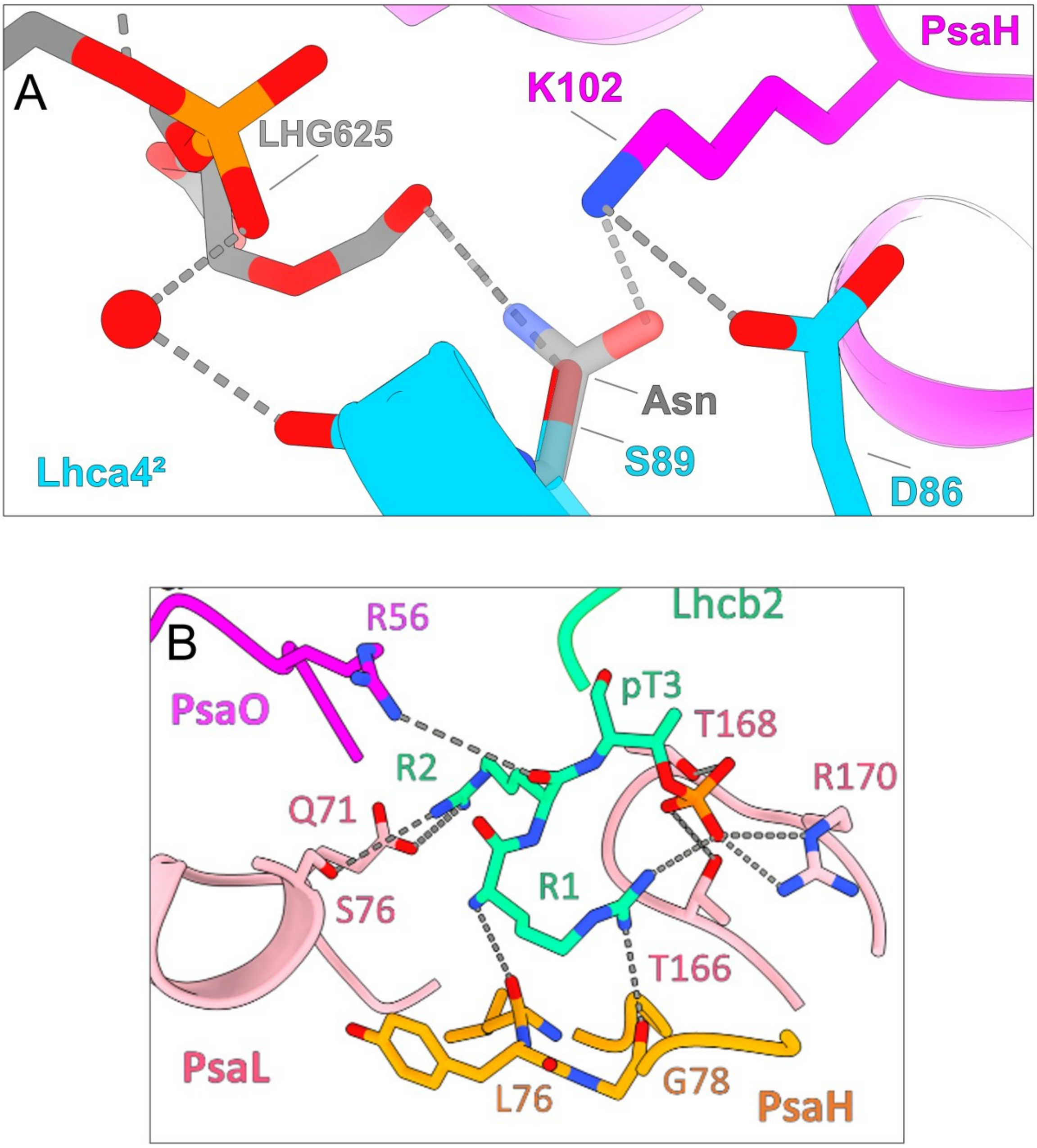

**Supplemental Figure 5. Interface features in the L-PSI-LHCII.**

**(A)** Model based mutagenesis of the Ans→Ser89 substitution in Lhca4. The atomic model of the domain connecting Lhca1 with PsaH in *P. oceanica* L-PSI-LHCII is presented. A rotamer of the substituting Asn and its potential interaction from land plants is shown in transparency. **(B)** Stromal view of the LHCII-Core interface highlighting the interactions of the Phosphorylated N-ter (P-Thr3)of Lhcb2 with the core.

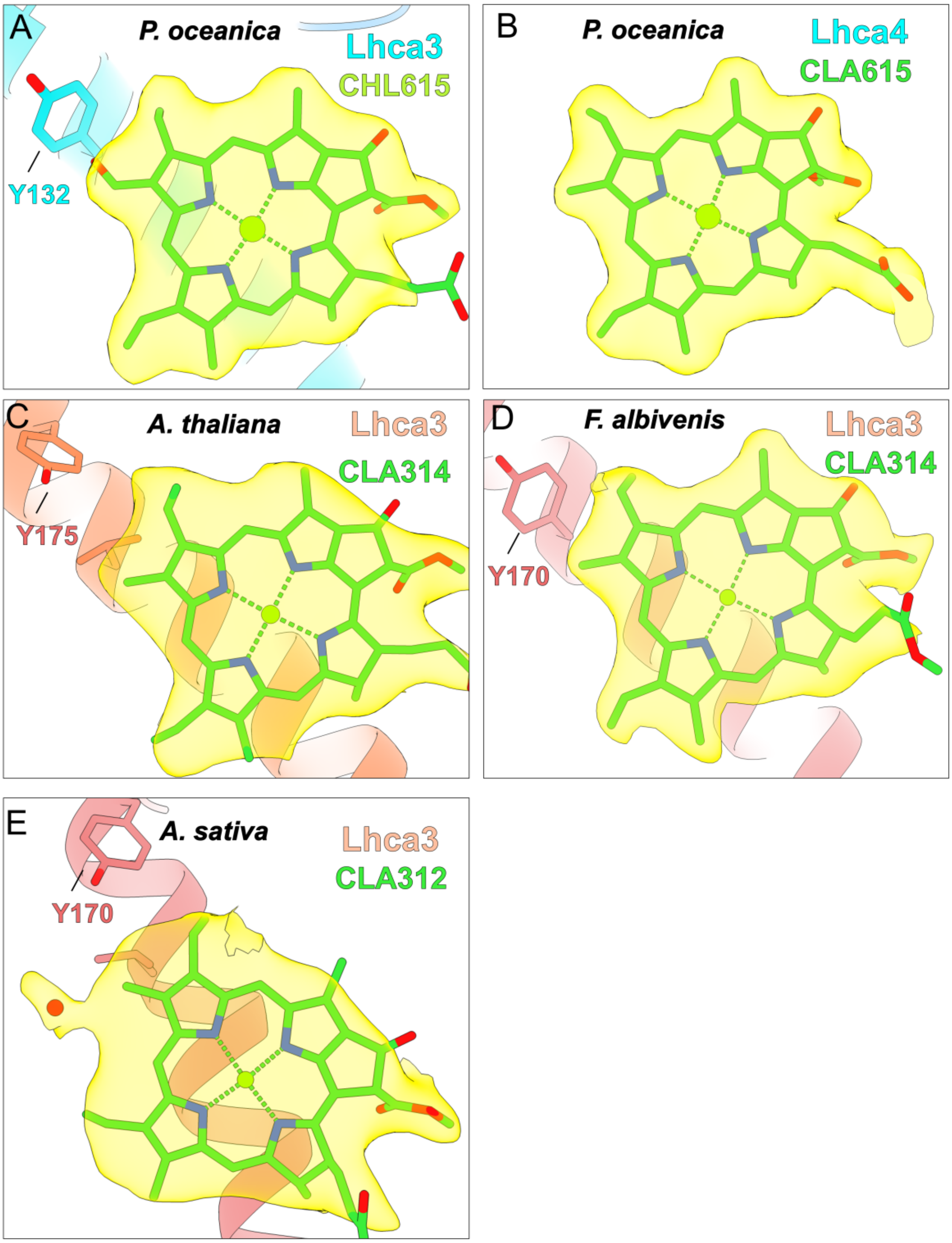

**Supplemental Figure 6. Atomic model and cryoEm density map of the Chl 615 sites and equivalents in the Lhca3 of representative species.**

**A.B.** Curent model and density map of *P. oceanica's* L-PSI-LHCII with a focus on CHL615 (A) and the Chl*a* occupying the equivalent position in Lhca4 (B). **C.** *A. thaliana*, PDB: 9GBI. **D.** *F. albivenis*, PDB: 8WGH. **E.** *A. sativa*, PDB: 8BCV. The Chls and residues numbers are shown as determined by authors. The Chls are shown as modelled by authors. The water molecule visible in E. was modelled based on the re-interpretation of the CryoEM map.

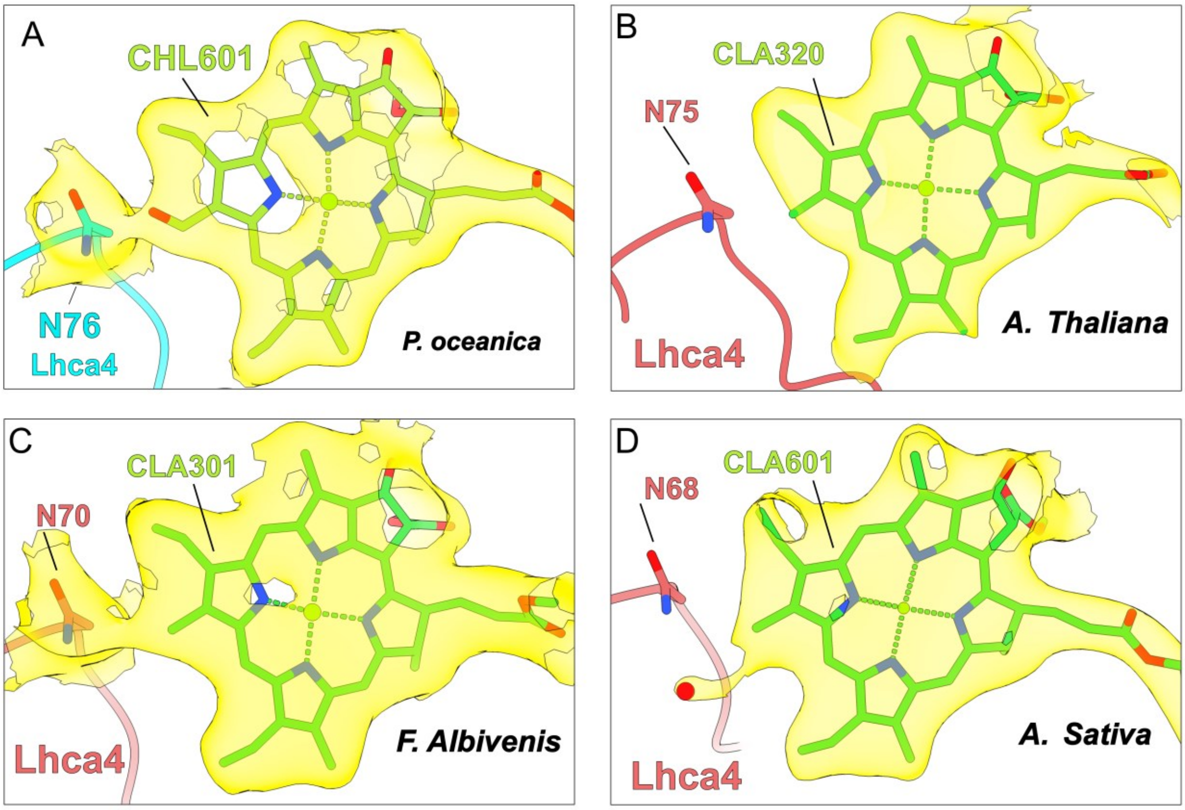

**Supplemental Figure 7. Atomic model and cryoEM density map of the Chl 601 and Asn76 sites and equivalent in the PSI of representative species.**

***A.*** *P. oceanica****.*** ***B.*** *A. thaliana*, PDB: 9GBI. **C.** *F. albivenis*, PDB: 8WGH. **D**. *A. sativa*, PDB: 8BCV. The Chls and residues numbers are shown as determined by authors. The Chls are shown as modelled by authors. The water molecule visible in C. was modelled based on the re-interpretation of the CryoEM map.

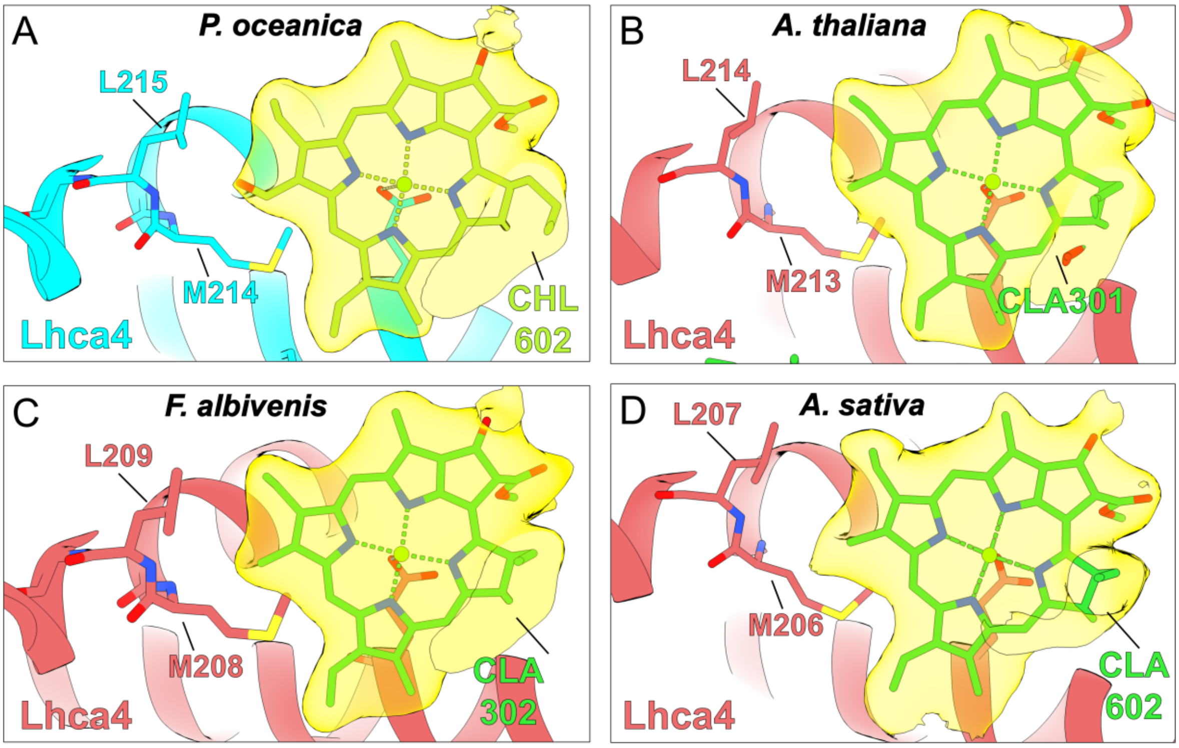

**Supplemental Figure 8. Atomic model and cryoEM density map of the Chl 602 sites and equivalent in the Lhca4 and LHCII of representative species.**
**A.** *P. oceanica*. **B.** *A. thaliana*, PDB: 9GBI. **C.** *F. albivenis*, PDB: 8WGH. **D**. *A. sativa*, PDB: 8BCV. The Chls and residues numbers are shown as determined by authors. The Chls are shown as modelled by authors.

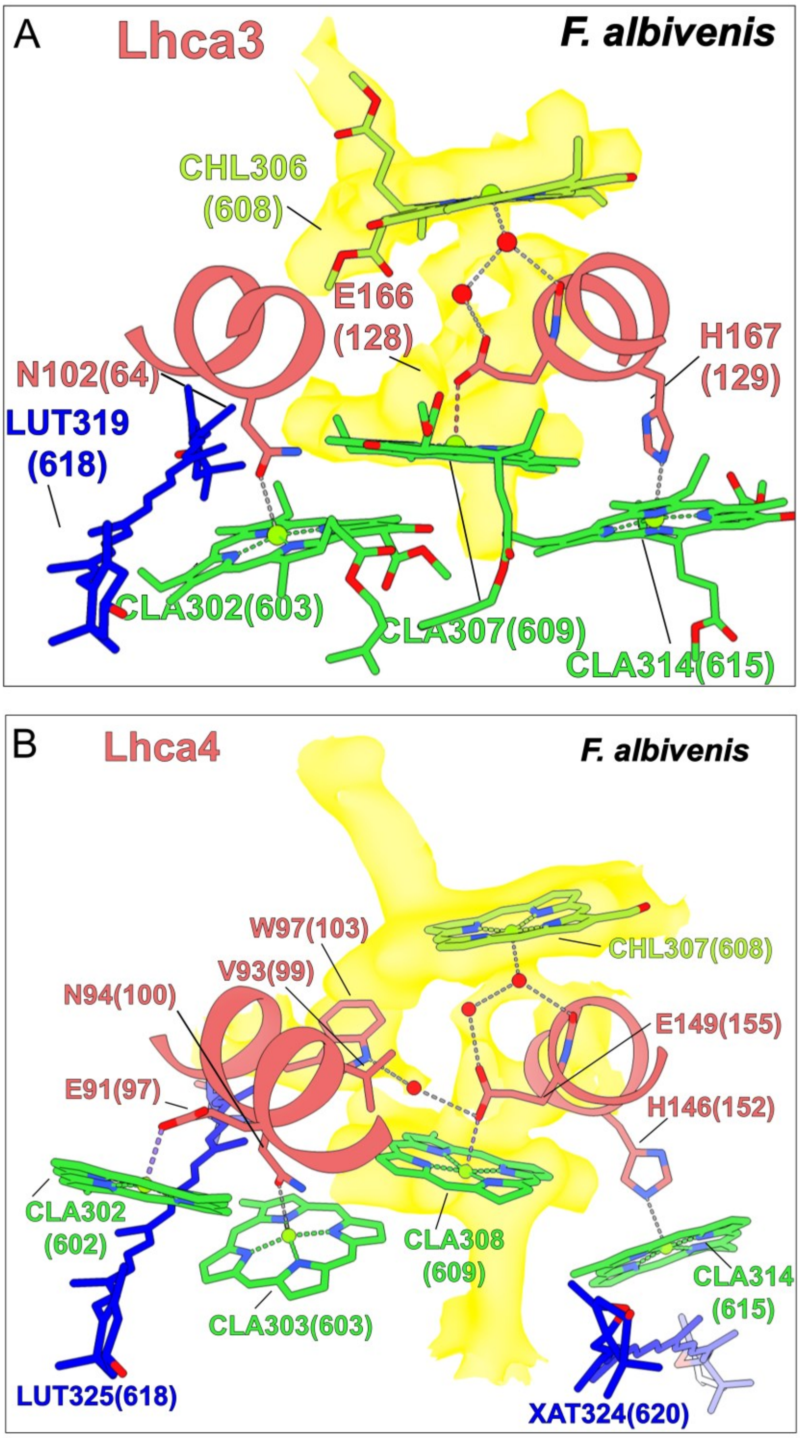

**Supplemental Figure 9. Overall organisation and pigments content of the far-red domain of Lhca3 and Lhca4 in *F. albivenis*.**

Architecture and co-factors composition of the far-red domain in Lhca3 (**A**) and Lhca4 (**B**) of *F. albivenis* (PDB: 8WGH). The Chls and residues numbers are shown as determined by authors with the equivalent position number in *P. oceanica* into brackets. Water molecules are represented by red spheres. CryoEM density is shown in yellow only for highlighting the residues and the water molecules network bridging CHL608 and CLA609. CryoEM maps were reinterpreted and the sites were remodelled to integrate the water molecules.

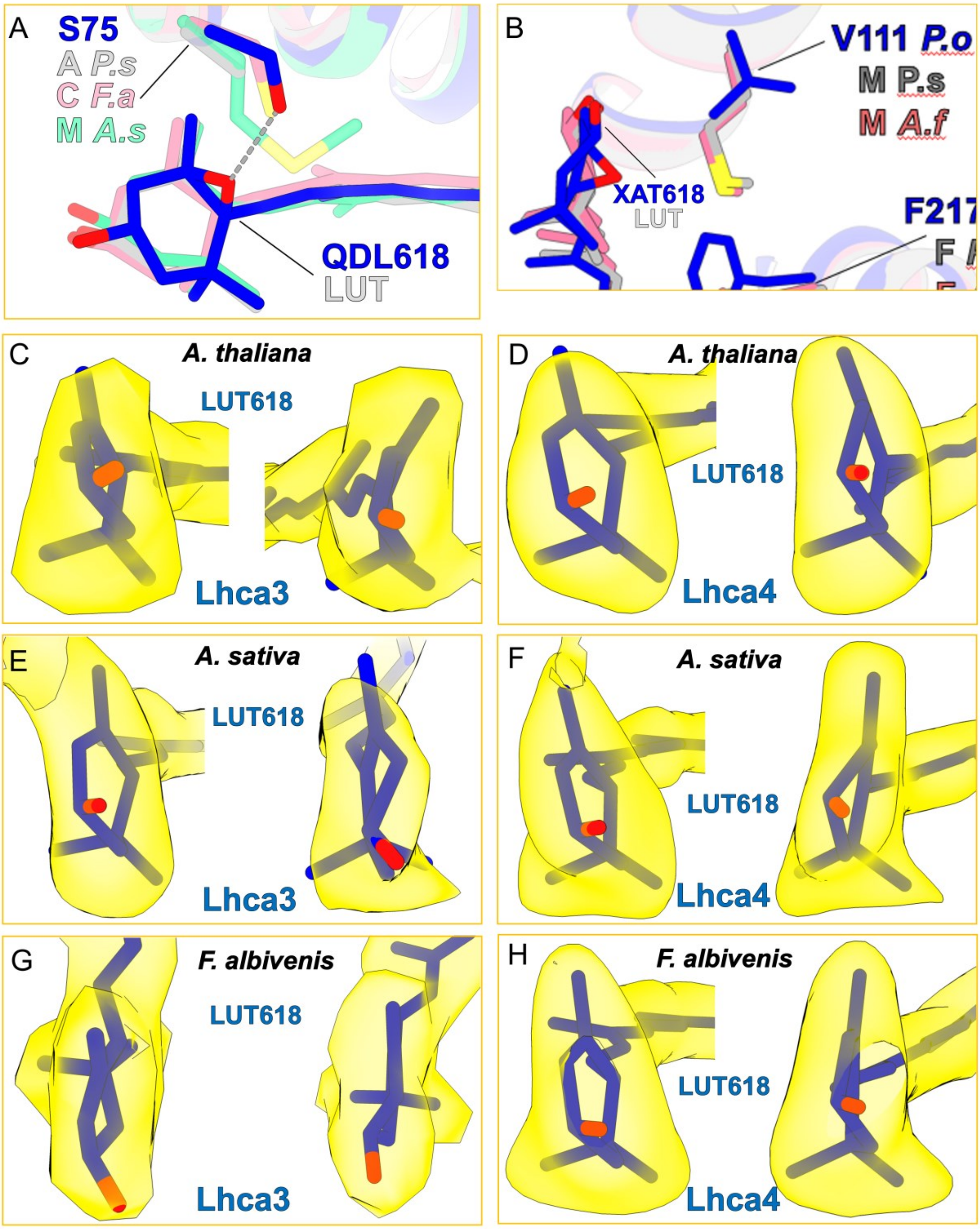

**Supplemental Figure 10. Atomic model and cryoEM density map of the lutein ionone rings at Carotenoid 618 sites and equivalent in the Lhca3 and Lhca4 of representative species. A.B.** Comparison of the protein environment and pigment organization around position 618 in Lhca3 (A) and Lhca4 (B) across different species. **C.D.** *A. thaliana*, PDB: 9GBI. **E.F.** *A. sativa*, PDB: 8BCV. **G.H.** *F. albivenis*, PDB: 8WGH. These positions were modeled as violaxanthin in *A. thaliana* (9GBI) and *A. sativa* (8BCV) and as lutein (Lhca3) and zeaxanthine (Lhca4) in *F. albivenis*. CryoEM map were reinterpretated and the sites were remodelled to integrate a lutein.

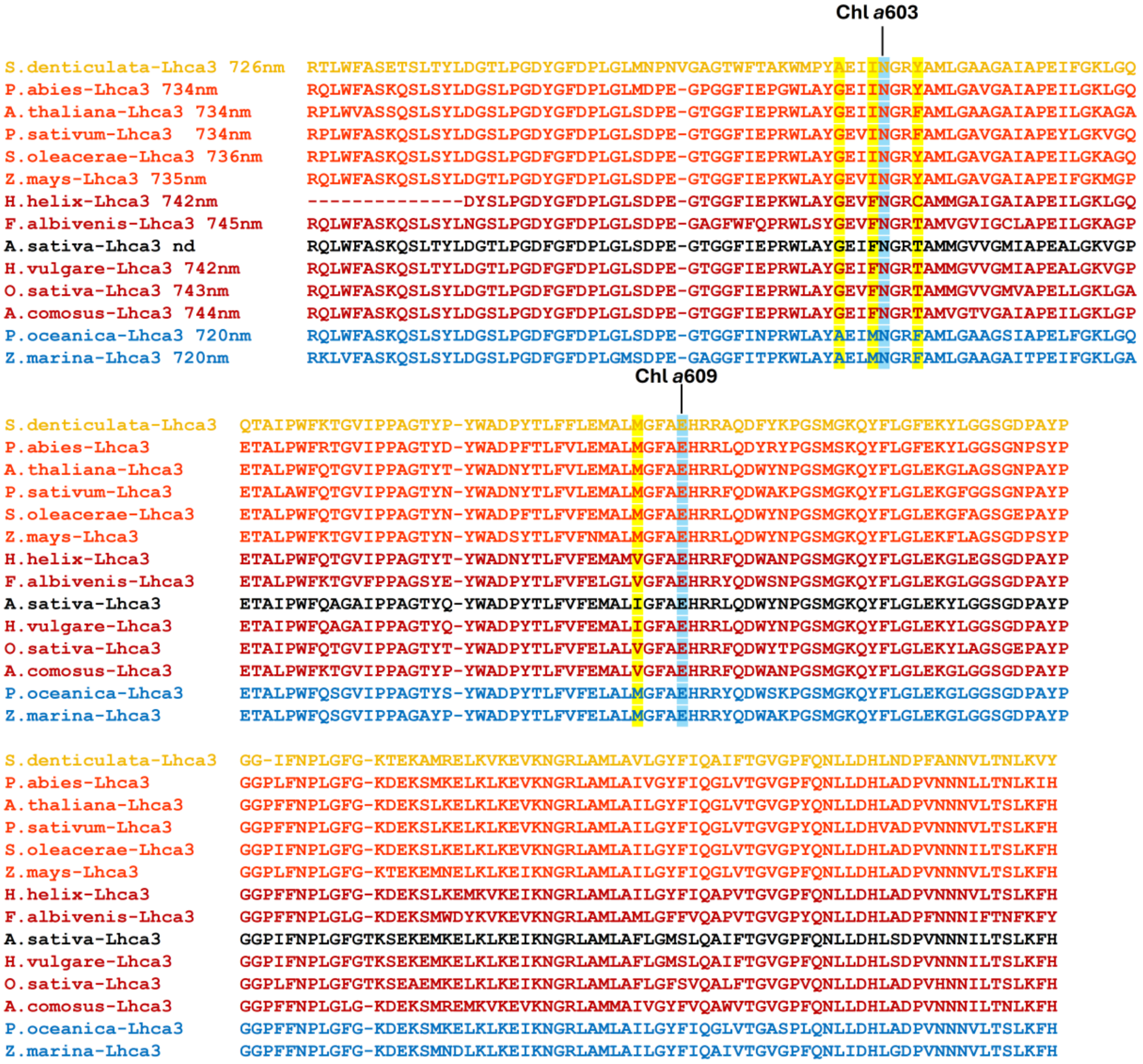

**Supplemental Figure 11. Sequence alignment of Lhca3 from representative species.** The species were colored and grouped based on the extent of PSI Far-red emission. Positions whose variation correlate with the main spectral windows across species are highlighted in yellow (see Fig. 4 and 5). Highlighted in blue are the co-axial ligands of Chl *a*603 (N) and Chl *a*609 (E).

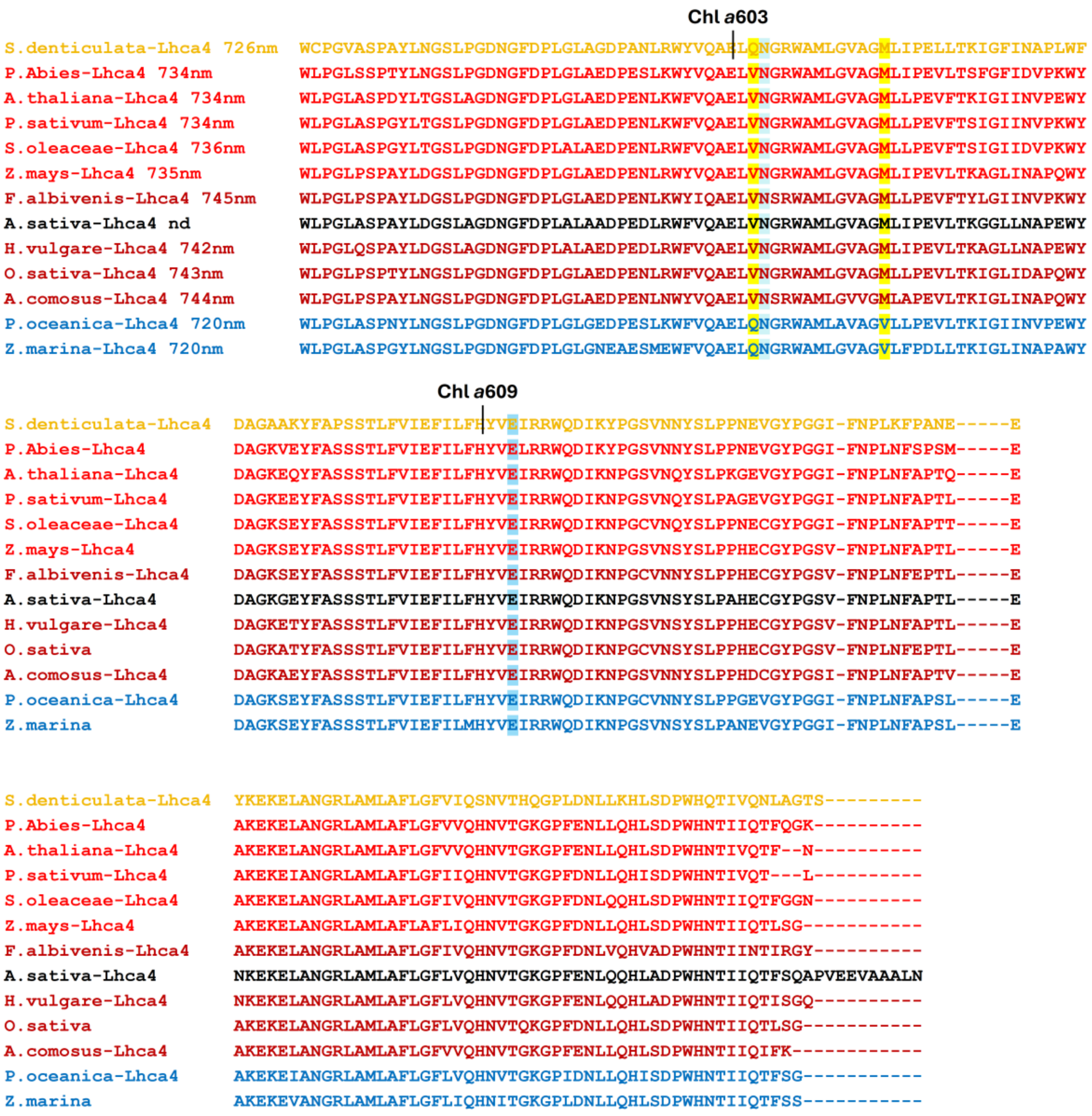

**Supplemental Figure 12. Sequence alignment of Lhca4 from representative species.**
The species were colored and grouped based on the extent of PSI Far-red emission. Positions whose variation correlate with the main spectral windows across species are highlighted in yellow (see Fig. 4 and 5). Highlighted in blue are the co-axial ligands of Chl *a*603 (N) and Chl *a*609 (E).

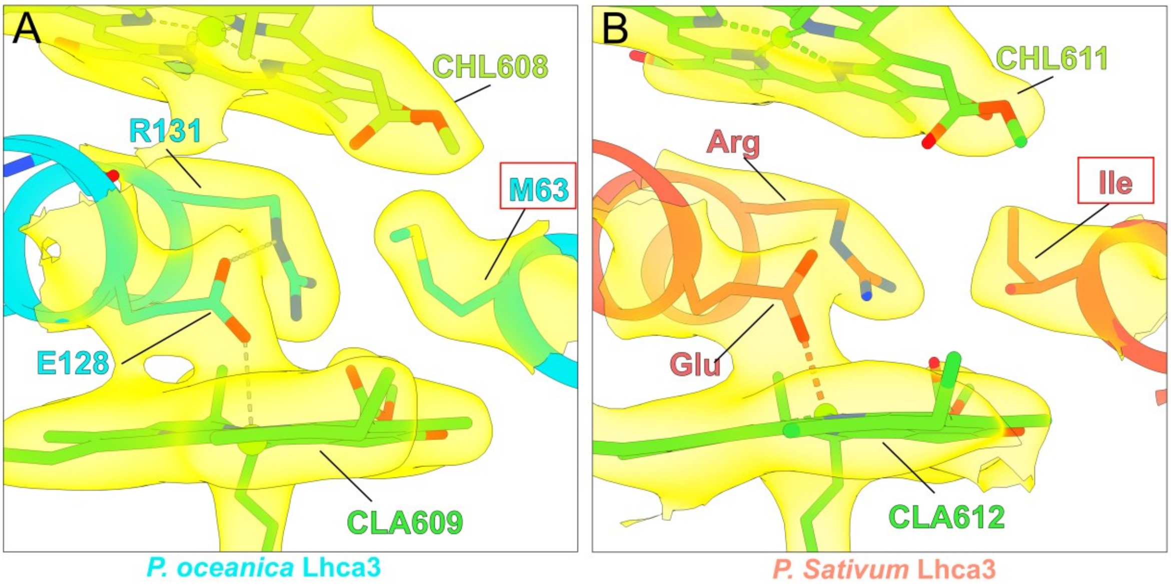

**Supplemental Figure 13. Atomic model and CryoEM density map around position 63 in the far-red domain of Lhca3 in *P. oceanica* and *P. sativum*.**

**A.B.** Overhaul organization of the far-red domain of Lhca3 in the vicinity of Val or Met 63 in *P. oceanica* (A) and *P. sativum* (B).

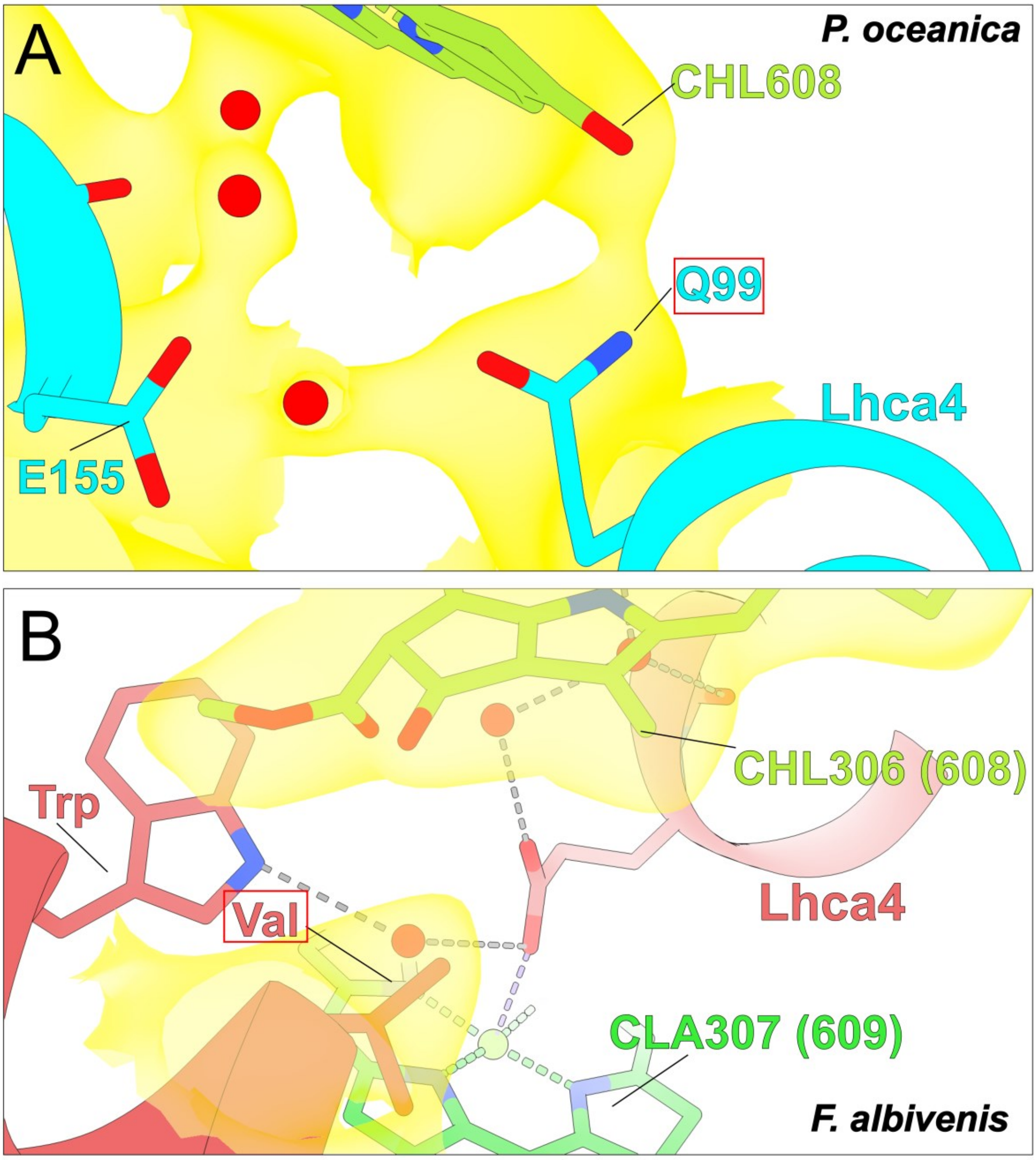

**Supplemental Figure 14. Comparison of residues at position 99 and in Lhca4 from *P. oceanica* and *F.albivenis*.**

**A.** Atomic model and CryoEM density map showing the interaction of Gln99 with CHL608 in *P. oceanica*'s Lhca4. **B.** Atomic model and CryoEM density map of *F. albivenis's* focused on Val (equivalent 99 in P. *oceanica*) and CHL306 (equivalent of CHL608). The Chls numbers are shown as determined by authors with the equivalent position number in *P. oceanica* into brakets.

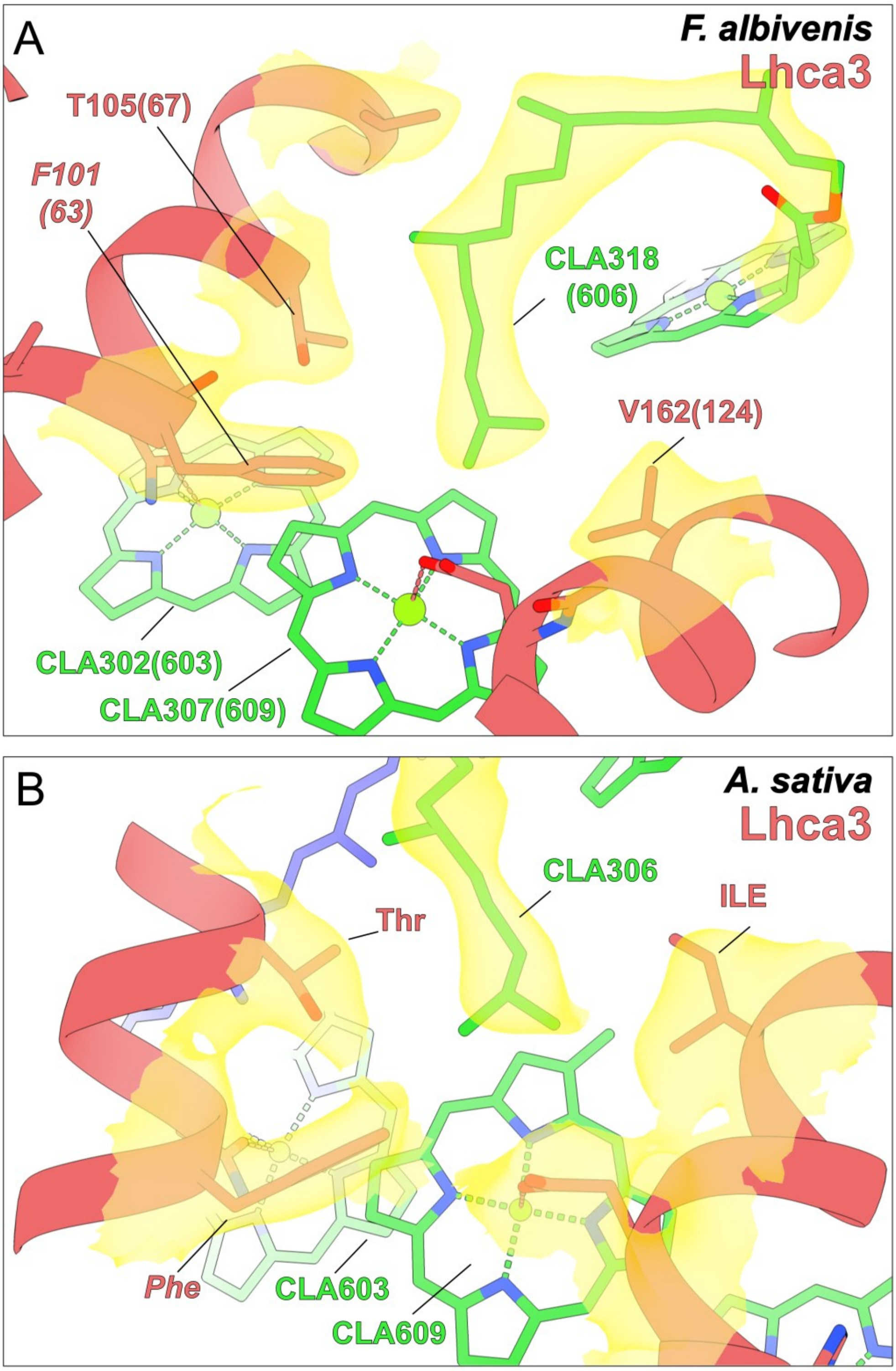

**Supplemental Figure 15. Atomic model and CryoEM density map of the residues surrounding Chl *a*609 in Lhca3 from representative species.**

**A.** *F. albivenis* (PDB: 8WGH). **B.** *A. sativa (*PDB: 8BCV). CryoEM maps were reinterpreted and the sites were remodelled to integrate the phytol chain of CLA606 (CLA318 in *F. albivenis*). The residues and Chls numbers are shown as determined by authors with the equivalent position number in *P. oceanica* into brackets.

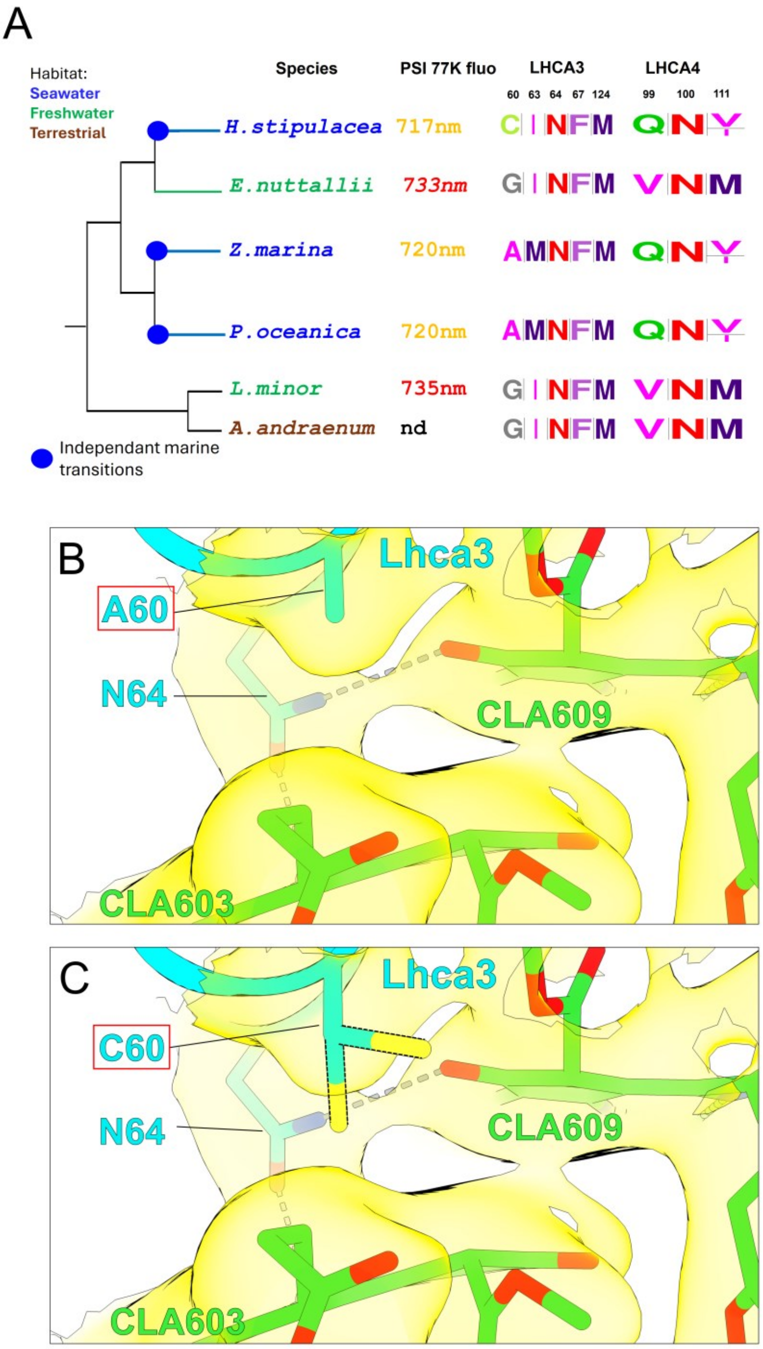

**Supplemental Figure 16. Diversification of the far-red domains in Alismatales representative species.**

**A.** Cladogramme of representative alismatales species with their respective 77k PSI emission and the amino acid occupying the position identified in Figure 4. The habitat of each taxa is represented by a different colour as indicated. **B**. Atomic model and cryo-EM density map of the residues surrounding CLA603 and CLA609 in *P. oceanica* Lhca3. **C.** Model based mutagenesis of the domain shows in panel B where the Ala63 was replaced by a Cys (two rotamers are shown) to predict the impact of the Ala-to-Cys substitution in *H. stipulacea* (panel A).

**Supplemental Table 1. Summary of the data collection, refinement, and validation statistics of the L-PSI–LHCII structure.**

| | **L-PSI–LHCII** (PDB: xxxx, EMDB: EMD-xxxx) |
|---|---|
| Data collection and processing | |
| Magnification | 165,000 |
| Voltage (kV) | 300 |
| Electron exposure (e–/Å2) | 43 |
| Defocus range (μm) | -0.5 to -1.9 |
| Pixel size (Å) | 0.729 |
| Initial particle images (no.) | 13,810 |
| Symmetry imposed | C1 |
| Final particle images (no.) | 73,859 |
| Map resolution (Å)<br>FSC threshold 0.143 | 1.87 |
| Refinement | (see below) |

Refinement

| **Stat** | **Dimer** | **Trimer** | **Core** |
|---|---|---|---|
| Clashscore, all atoms | 5.90 | 3.02 | 0.59 |
| Poor rotamers | 0.21% | 0.13% | 0.40% |
| Favored rotamers | 88.87% | 86.99% | 90.29% |
| Ramachandran outliers | 0 | 1 | 1 |
| Ramachandran favored | 98.80% | 98.17% | 98.06% |
| Rama distribution Z-score | 2.70 ± 0.36 | 1.67 ± 0.26 | 1.76 ± 0.15 |
| MolProbity score | 1.32 | 1.09 | 0.70 |
| Cβ deviations > 0.25 Å | 0.00% | 0.00% | 0.00% |
| Bad bonds | 1.13% | 1.15% | 1.35% |
| Bad angles | 0.17% | 0.09% | 0.17% |
| Cis Prolines | 0.00% | 0.00% | 1.01% |
| CABLAM outliers | 0.9% | 0.9% | 1.0% |
| CA geometry outliers | 0.00% | 0.53% | 0.33% |

| | |
|---|---|
| Initial model used (PDB code) | xxx |
| Map sharpening B factor (Å2) | -5.1 |

| | |
|---|---|
| Model composition | |
| Non-hydrogen atoms | 16,679 |
| Protein residues | 1,399 |
| Ligands | 129 |
| Waters | 285 |
| B factors (Å2) (min/max/mean) | |
| Protein | 16.21/119.59/43.34 |
| Ligand | 16.34/123.46/48.16 |
| Waters | 22.45/91.18/43.86 |
| R.m.s. deviations | |
| Bond lengths (Å) | 0.009 |
| Bond angles (°) | 1.088 |
| Validation | |
| MolProbity score | 1.33 |
| Clashscore | 6.00 |
| Poor rotamers (%) | 0.27 |
| Ramachandran plot | |
| Favored (%) | 98.18 |
| Allowed (%) | 1.82 |
| Disallowed (%) | 0.00 |

**Supplemental Table 2 : Excitonic coupling within the far-red pigment cluster of Lhca3 and Lhca4.**

Absolute excitonic couplings (|EC|) within the far-red pigment cluster of Lhca3 and Lhca4 in *P. oceanica* and *F. albivenis*. Values were calculated using a point-dipole approximation and are shown for inter-chlorophyll and chlorophyll–carotenoid pairs.

| Paralogs | Coupling ($cm^{-1}$) | *P. oceanica* | *F. albivenis* | Δ P. o vs F. a |
|---|---|---|---|---|
| Lhca3 | \|EC(603-609)\| | 187 | 213 | -12 % |
| | \|EC(603-Car)\| | 582 | 481 | +21 % |
| Lhca4 | \|EC(603-609)\| | 187 | 228 | -18 % |
| | \|EC(603-Car)\| | 645 | 573 | +13 % |
